# Body mass index trajectories preceding first report of poor self-rated health: a longitudinal case-control analysis of the English Longitudinal Study of Ageing

**DOI:** 10.1101/405563

**Authors:** Adam Hulman, Daniel B Ibsen, Anne Sofie D Laursen, Christina C Dahm

## Abstract

**Background:** Studies have consistently found that obesity is associated with poor self-rated health, but how body mass index (BMI) developed in the lead up to poor self-rated health is unknown.

**Methods:** We nested a longitudinal case-control study in the English Longitudinal Study of Ageing (1998-2015) to investigate BMI trajectories in the years preceding a first self-report of poor health. Participants rated their health at each data collection; every other collection included a BMI assessment by a nurse. Case status was defined as a first report of poor health during follow-up. Three age- and sex-matched controls were identified per case using density sampling. BMI trajectories were fitted to time backwards prior to first report of poor health using mixed-effects models. Age and sex were potential modifiers. We conducted subgroup analyses of those not reporting certain chronic diseases or smoking.

**Results:** We identified 732 cases and 2195 controls. Age, but not sex, modified the association between BMI and self-rated health. Participants reporting poor health at age 60 had a 1.5 kg/m^2^ (95%CI: 0.8 to 2.1) higher BMI at the time of reporting than controls, and their BMI had previously increased sharply. After age 75, cases no longer had higher BMI than controls, and their BMI had decreased sharply prior to reporting poor health. Age was also an effect modifier among those without diabetes, however BMI trajectories were more similar among the middle-aged.

**Conclusion:** Development of BMI was associated with poor self-rated health; however, the nature of the association depended markedly on age.

What is already known on this subject

- Studies based on a single body mass index measurement have consistently found that being underweight or obese was associated with poor self-rated health.
- How body mass index developed in the lead up to poor self-rated health was unknown.

What this study adds

- In the present study, body mass index trajectories leading up to poor self-rated health were strongly dependent on age at first report of poor health. Middle-aged participants with poor self-rated health had higher body mass index than controls; both groups had increasing body mass index over time.
- In contrast, a report of poor health at age 75 or later was preceded by a decrease in body mass index; controls were 10 years older than cases before they started to lose body mass index.
- Our results suggest that future prospective studies and clinical decision making should take into account the age-dependent association between development of body mass index preceding poor self-rated health.

## INTRODUCTION

Self-rated health is a robust and independent predictor of health outcomes and mortality.[1-2] It is a construct that encapsulates physical, mental and social dimensions of health. The weighting of these dimensions within each individual is generally unknown,[3] although studies indicate that self-assessment of physical functioning is a substantial contributor to self-rated overall health.[4-5] Observational studies have investigated whether body mass index (BMI) is associated with self-rated health among middle-aged adults, generally finding that obese persons are more likely to report poor health and to live more years with poor health than persons of normal weight.[6-8]

The mechanisms underlying the association between BMI and self-rated health are likely to vary substantially according to age, as conditions contributing to self-rated health change in importance across the lifespan. Developments in BMI may be important, because in sedentary populations BMI mainly reflects fat mass,[9] which may affect physical functioning and incidence of disease such as type 2 diabetes. However, among the elderly, changes in BMI generally reflect loss of muscle mass, and sarcopenia has been associated with greater mortality.[10] Chronic conditions, such as chronic obstructive pulmonary disease (COPD), are among the main potential causes of unintended weight loss among elderly.[11] Despite the health implications of different body compositions at a given BMI, measurement of BMI remains the most common assessment of adiposity in the general population. Yet, little is known about the development of BMI prior to poor self-rated health, nor how this may differ by age and sex.

We investigated differences in BMI trajectories up to 10 years prior to a first report of poor self-rated heath, compared to age- and sex-matched controls, taking into account a potential modifying effect of age at first report of poor self-rated health, and sex. We hypothesized that poor self-rated health in middle-aged individuals would be positively associated with higher BMI in preceding years, but that this association would be weaker at older ages.

## METHODS

### Study population

Our study is based on data from the English Longitudinal Study of Ageing (ELSA).[12] ELSA participants were sampled from three years of the Health Survey for England (1998, 1999 and 2001) that we consider as wave 0 in the current study. Only those born before 1 March 1952 were considered as core members as the goal of the original study was to recruit a sample representing the population older than 50 years. Participants were followed-up and interviewed every two-three years between 1998 and 2015 (waves 0-7). Even waves included a nurse visit with clinical measurements. Informed consent was obtained from all participants and the study was conducted according to the Declaration of Helsinki.

We designed a longitudinal case-control study to be able to compare differences in BMI trajectories between those reporting poor health for the first time during follow-up and the source population not reporting poor health. The selection of cases is described in Figure 1. As we were examining BMI trajectories before first self-report of poor health, we excluded those who already had reported poor health at waves 0-1 and those with fewer than two BMI measurements. To compare BMI trajectories with a control sample not reporting poor health up to and including the wave of the index case, we selected three controls per case using density sampling, matching on age (±1 year) and sex. The wave in which an index case reported poor health defined the time of event occurrence and defined time=0 for the controls.

**Figure 1.**
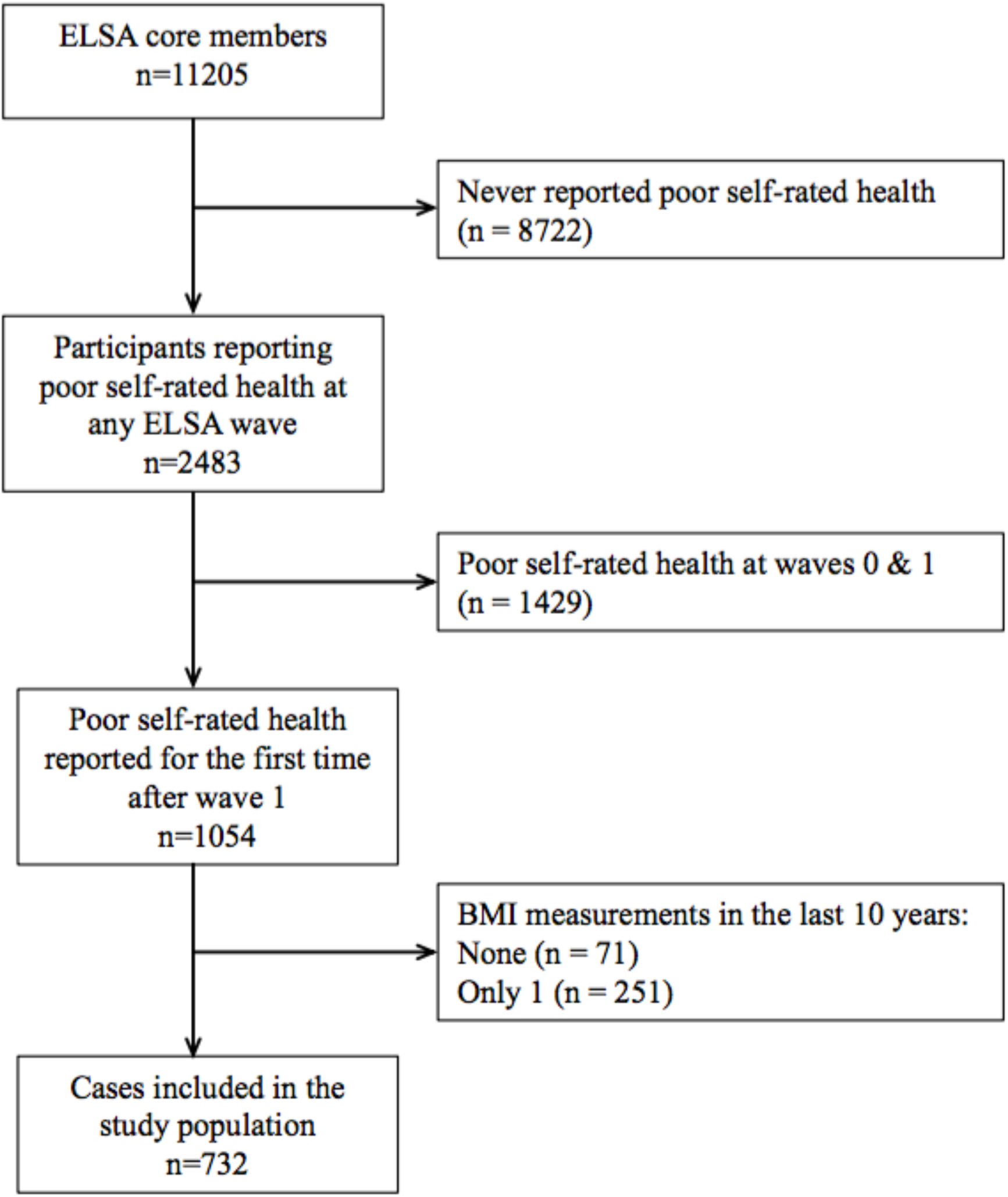
Identification of cases.

### Self-rated health

Self-rated health was assessed in each wave during an interview using the following question: “Would you say your health is (1) excellent, (2) very good, (3) good, (4) fair or (5) poor?” Our outcome was defined as answering (5) poor. The question was slightly different in waves 0 and 3, where the answer (1) excellent was dropped and (5) poor was replaced with “bad” and “very bad”. We combined these two to represent poor health.

### Body mass index

BMI was calculated using weight and height measurements performed by trained personnel during nurse visits or clinical examinations (BMI (kg/m^2^)=weight/height-squared). The measurements were conducted at even waves. If a participant was chair-bound then an estimate was obtained from the respondent instead. If the nurse thought that these were likely to deviate from the true figures more than 2 cm for height and 1 kg for weight, then they were marked as unreliable and were not used in our analysis.

### Covariates

Information on socioeconomic class, smoking and marital status was assessed with standard questionnaires. Smoking status was defined as a binary variable (never vs. ever) and was used as a time-invariant factor assuming that people are not likely to start smoking in this age if they never smoked before. Depression was assessed with the eight-item version of the Center for Epidemiologic Studies Depression Scale (CES-D) at each wave. Diabetes, cardiovascular disease (heart attack or stroke), cancer and chronic lung disease status was assessed and updated at each wave based on self-reports.

### Statistical analysis

BMI trajectories were fitted using mixed-effects models including both a random intercept and a slope. This approach accounts for the repeated measurement structure in the data set. Models were specified with a linear term for time. Time was defined as years backwards from first report of poor health (event outcome) among cases and a corresponding matched time for controls. We considered BMI measurements for up to ten years before the event. A binary indicator variable was defined to distinguish cases from controls. We included this variable and its interaction with time in the model to assess differences in both level (intercept) and slope, respectively, between cases and controls. Then we included age at event occurrence (time=0) and its interaction with time in the model, to investigate the effect of age on the level and slope of the trajectories. We also included the interaction between the case-indicator variable and age to examine whether age had a differential effect on the level of BMI between cases and controls. Finally, we included a three-way interaction between the case-indicator, age and time, to examine whether age had a differential effect on the BMI slope between cases and controls. We calculated how much of the between-person variation of the intercepts and the slopes was explained by age. We also tested whether sex had a modifying effect on BMI development by including the main term and an interaction with time in a model. The final model was fitted with and without adjustment for socioeconomic class. We further conducted five subgroup analyses including only participants without diabetes, cardiovascular disease, cancer, chronic lung disease or never-smokers at baseline, as these factors are associated with BMI changes.[13-16]

Statistical analyses were conducted using the nlme (v3.1-128) and Epi (2.9) packages in R (3.3.1). Full documentation is available in the online supplementary appendix.

## RESULTS

Almost 80% of the 11,205 core members did not report poor health at any point in the study, leaving 2483 potential cases (Figure 1). After excluding those who reported poor health at waves 0 or 1 (n=1429), those with zero (n=71) or only one (n=251) BMI measurement, the final case sample included 732 individuals (56% women). Three controls were successfully matched to all but one case, where only two controls were identified, resulting in a final study population of 732 cases and 2195 controls.

A quarter of cases were younger than 64 years of age when they reported poor health, while a quarter were older than 80 years (Table 1). Compared to controls, cases were more likely to be in a lower socioeconomic class, divorced or separated at baseline, and a lower proportion reported that they had never smoked. Cases were more likely to have more depression symptoms and to have diabetes, cardiovascular disease or cancer than controls. Cases and controls had on average 2.3 and 2.4 BMI measurements, respectively.

**Table 1:** Characteristics of the study population at first report of poor health, the English Longitudinal Study of Ageing, UK, 1998-2015.

|  | Case | Control |
| --- | --- | --- |
| N | 732 | 2195 |
| Age (year)<sup>1</sup> | 73 (64-80) | 72 (64-80) |
| Sex (male, %) | 44 | 44 |
| Social class (%)<sup>2,3</sup> |  |  |
| Professional or managerial technical | 31 | 38 |
| Skilled non-manual | 22 | 26 |
| Skilled manual | 18 | 17 |
| Semi-skilled or unskilled manual | 29 | 19 |
| Marital status (%)<sup>2</sup> |  |  |
| Single | 5 | 5 |
| Married | 61 | 67 |
| Divorced/separated | 14 | 9 |
| Widowed | 21 | 19 |
| Never smoked (%)<sup>2</sup> | 29 | 38 |
| Depression score (0-8)<sup>4</sup> | 1 (0-2) | 3 (1-5) |
| Diabetes (%) | 19 | 9 |
| Cardiovascular disease (%) | 21 | 11 |
| Cancer (%) | 21 | 6 |
| Chronic lung disease (%) | 15 | 5 |
| Self-rated health (%) |  |  |
| Excellent | 0 | 15 |
| Very good | 0 | 34 |
| Good | 0 | 35 |
| Fair | 0 | 17 |
| Poor | 100 | 0 |
| Number of BMI measurements (%)<sup>5,6</sup> |  |  |
| 2 | 70 | 64 |
| 3 | 30 | 36 |

<sup>1</sup>median age at first report of poor self-rated health (cases) or age when sampled (controls) and interquartile range
<sup>2</sup>information assessed at wave 1
<sup>3</sup>missing for 3% of the cohort (n=83)
<sup>4</sup>Center for Epidemiologic Studies Depression Scale (CES-D) ranging from 0 to 8
<sup>5</sup>within the last 10 years
<sup>6</sup>one control had four measurement

Age had a modifying effect on the association between BMI trajectories and self-rated health, but we found no sex differences in BMI trajectories. Estimated BMI levels ten years before and at first report of poor self-rated health, and BMI slopes by age at reporting poor self-rated health, are presented for cases and controls in Figure 2. Modelled BMI trajectories by self-reported health for four different ages at first report of poor health are presented in Figure 3A. The average case and control were overweight (25-30 kg/m^2^) throughout the ten years, regardless of age. Ten years before first report of poor health, cases had higher BMI than controls, regardless of age (Figure 2A; see online supplementary appendix, page 39). However, differences in BMI at first report of poor health depended strongly on age (Figure 2B). Poor self-rated health was associated with a higher BMI at first report of poor health before 76 years of age but no difference afterwards. E.g. participants reporting poor health at age 60 had a 1.5 kg/m^2^ (95%CI: 0.8 to 2.1) higher BMI at first report of poor health than controls. Slopes were similarly increasing among middle-aged cases and controls, but cases reporting poor health at age 75 and later exhibited a declining pattern in BMI preceding their report of poor health. Corresponding controls were on more stable trajectories, such decline in BMI happened at an approx. 10 years older age. The difference in slopes between the two groups increased with age (casexage(10y)xtime(10y) interaction: -0.6 95%CI: -0.9 to - 0.3; p=0.0004). Adding age at first report of poor health and its interaction with time to the model explained a substantial portion (13%) of the between-person variation in slopes, but not in levels of BMI (2%). Adjustment for socioeconomic class did not have a major influence on our results (see online supplementary appendix, pages 36-38). There were even larger level differences between cases and controls among participants who never smoked (n=1047; 36%) and the effect of age at first report of poor health on BMI slopes was still present (Figure 3B). Cases and controls without diabetes (n=2576; 88%) were more similar to each other with respect to BMI in middle-age, but the divergence of slopes with ageing was still strongly present (casexage(10y)xtime(10y) interaction: -0.6 95%CI: -1.0 to -0.3; p=0.0003; Figure 3C). The subgroup analysis of those without cardiovascular disease (n=2532; 87%), cancer (n=2652; 91%) and chronic lung disease (n=2702; 92%) showed similar results to the main findings (Figure 3D-E-F).

**Figure 2.**
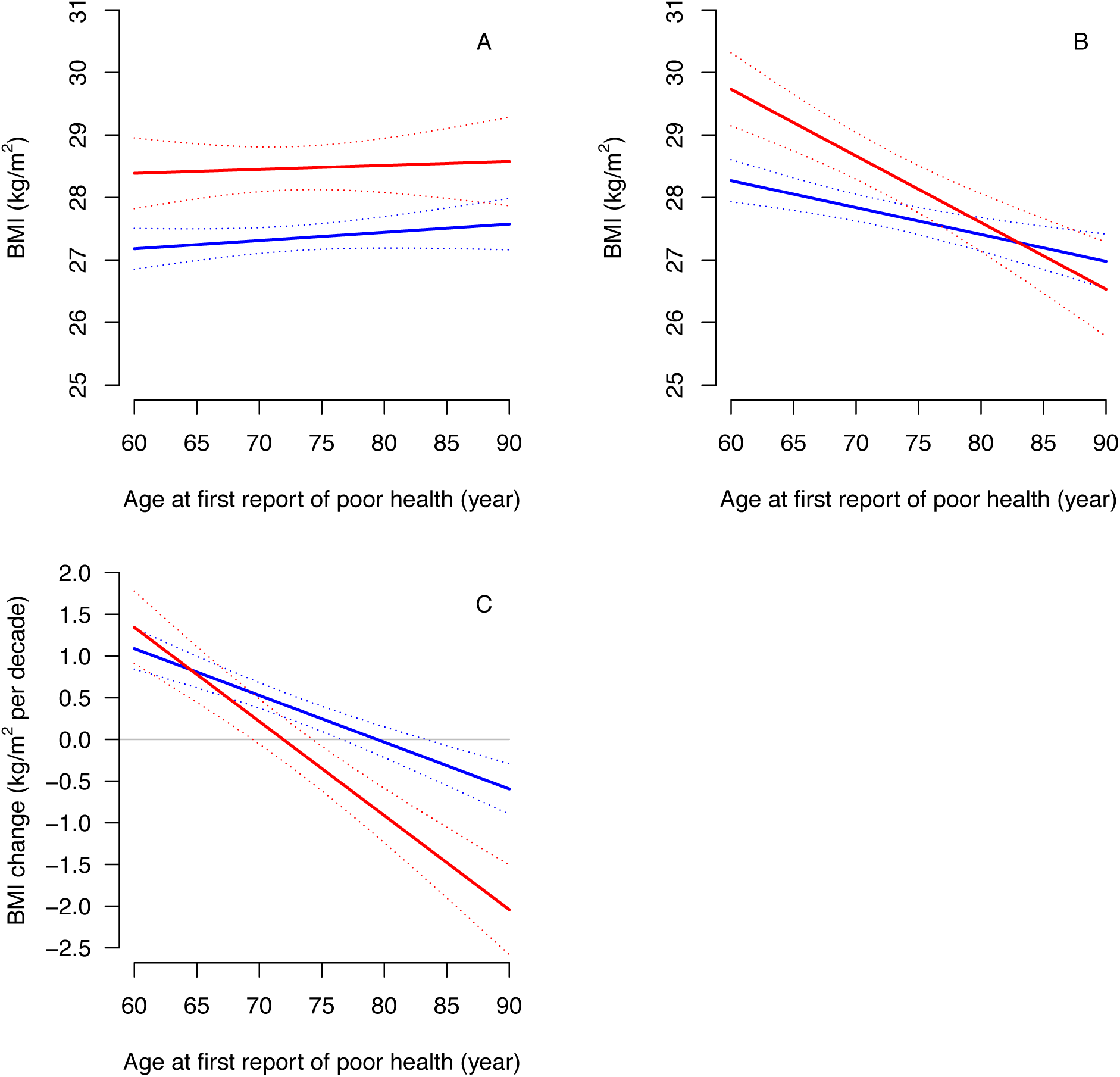
Estimated body mass index levels ten years before (A) and at first report of poor health (B) and BMI slopes (C) by age at first report of poor health (cases) or age when sampled (controls).

**Figure 3.**
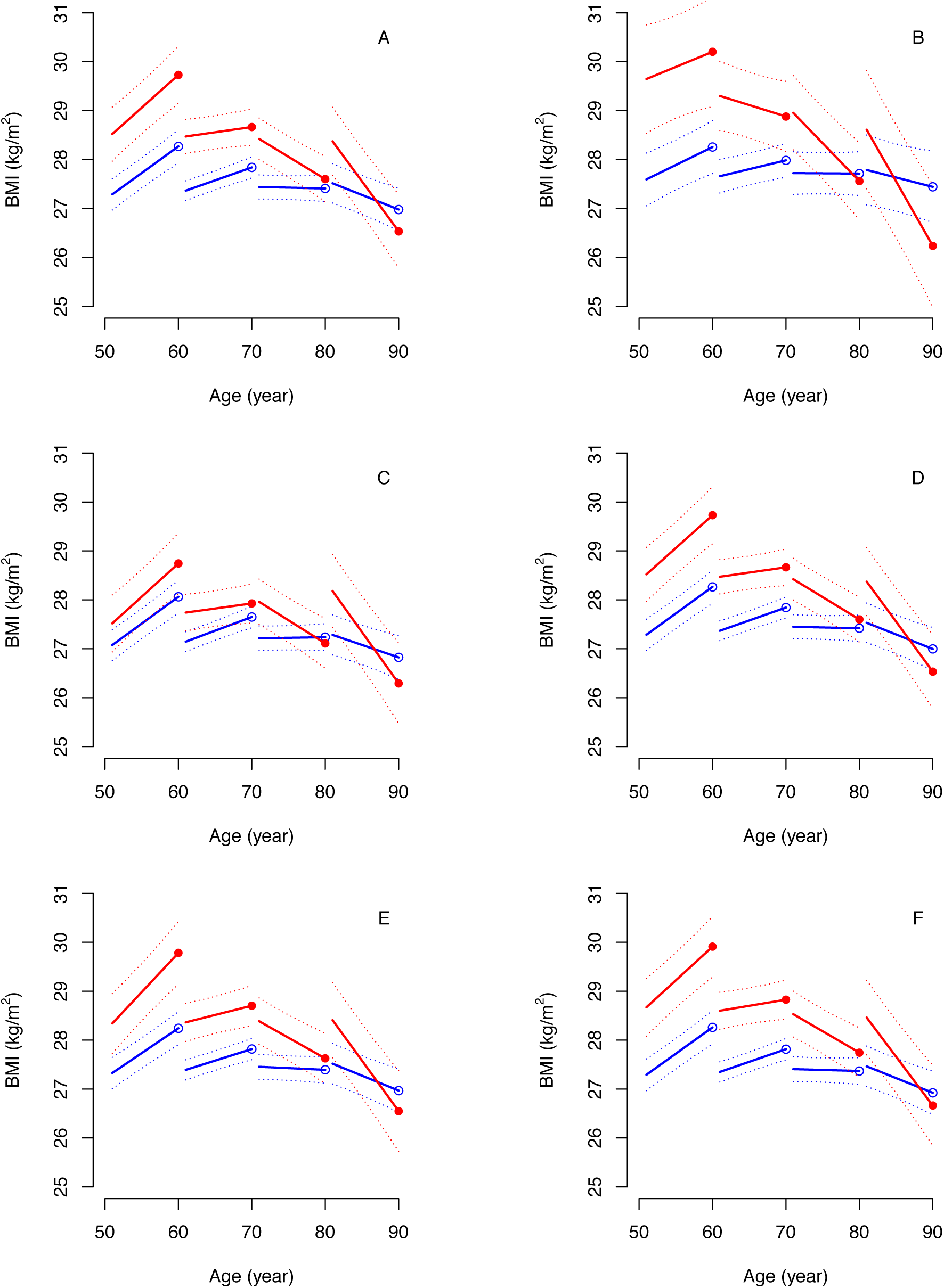
Modelled body mass index trajectories before first report of poor health at different ages (indicated by circles at 60, 70, 80 and 90 years) based on the entire sample (A) and those who never smoked (B), participants without diabetes (C), cardiovascular disease (D), cancer (E) and chronic lung disease (F). Cases: filled circle and red line; controls: empty circle and blue line.

## DISCUSSION

In this study of BMI trajectories prior to a first report of poor health in a middle-aged and elderly, generally overweight population, we found that development of BMI differed markedly between those who went on to report poor health, and their age- and sex-matched controls. These differences in trajectories varied by age, but were similar among men and women. Our results further indicate that among middle-aged men and women with poor self-rated health, a higher BMI is likely to be related to diabetes. Among the elderly, BMI decreased sharply prior to a report of poor health, regardless of diabetes, chronic lung disease and smoking status.

A major strength of our study is the use of multiple measurements of BMI performed by trained personnel, which enabled us to investigate trajectories of BMI prior to reporting poor self-rated health. Selection of controls using density sampling and matching on age and sex, provides a control group that was representative of the BMI trajectories experienced by participants in ELSA. Study participants had to have at least two valid BMI measurements by design, which led to an exclusion of 322 potential cases from the analyses. This missingness might have been related to both BMI and self-rated health, which could bias our estimates. We restricted our analysis to individuals who first rated their health poor during follow-up but not in waves 0 or 1, excluding 1429 participants. Therefore, our results may be generalizable mainly to people of good health until middle age. Lastly, given the observational nature of the study, the possibility of confounding cannot be excluded. For example, smoking has an effect on body weight and composition.[14] If smoking cessation is a consequence of a disease diagnosis leading to poor health (e.g. a cardiovascular event), resulting weight gain could have led to a less steep decline in BMI among elderly cases. However, we observed similar results in a subgroup who never smoked, which supports that our findings are not likely to be confounded by smoking.

Previous prospective studies of BMI and self-rated health have only investigated baseline BMI in relation to self-rated health during follow-up.[17-19] These studies have consistently found that a higher BMI at baseline was associated with a higher risk of poor self-rated health. Only few other studies have investigated changes in BMI in relation to future poor self-rated health, in very specific study populations.[20-21] Dennerstein et al. found concurrent weight gain to be associated with a decline in self-rated health among women during their menopausal transition.[20] In a study of female nurses older than 44 years, a shift from being underweight to being normal weight during a six-year period was associated with lower odds of experiencing worse self-rated health during follow-up, while a shift from normal weight to overweight was associated with higher odds of worse self-rated health.[21] The results from the studies of baseline BMI[11-13] and the studies of changes in BMI[14-15] are only in line with our results for middle-aged participants, but not the elderly. Analogous to these studies, a meta-analysis of 230 studies found a J-shaped dose-response association between BMI and all-cause mortality.[22] In an age-stratified analysis, this association was stronger among those aged <65 years old. In participants aged >65 years old, the strong association between BMI and mortality attenuated markedly in the overweight and obese range, whereas the association in the underweight range persisted. In our study, we also found a modifying effect of age, but here on the association between BMI trajectories and first report of poor self-rated health. Thus, future studies would benefit from taking this dynamically changing association across the lifespan into account.

Our findings suggest that there could be different etiologies underlying the BMI trajectory preceding poor self-rated health, depending on age. Obesity is linked to the development of many chronic diseases that have a major effect on general health, such as diabetes.[23] After restricting our analysis to individuals without diabetes, middle-aged, but not elderly, cases and controls had more similar BMI trajectories preceding poor self-rated health. Indeed, these results indicate that development of diabetes may partly explain the difference observed between middle-aged cases and controls. In contrast, among the elderly, a declining BMI trajectory preceding poor self-rated health was still present. COPD is one of the main causes of unintended weight loss among the elderly.[11] However, after restricting our analysis to participants without chronic lung disease or those who never smoked, we found very similar results to our main findings, suggesting that pulmonary diseases themselves cannot explain the differences in BMI slopes among the elderly. We had similar findings also with regard to cardiovascular disease and cancer. Another potential underlying mechanism explaining weight loss among the elderly could be depression related to the loss and bereavement of a spouse.[24] However, a higher proportion of controls were married at baseline, so it is not likely that cases experienced such an event more frequently during follow-up than controls, even though cases had more depression symptoms than controls.

Obesity is known to be associated with shorter healthy life expectancy.[6-8] In light of our results, tackling obesity earlier during the life course is crucial to achieve healthy ageing. Clinical practitioners and policy makers should consider age when suggesting weight loss to patients or planning interventions targeting weight loss. Given that self-rated health is a valid measure of experienced health, and is strongly related to objectively measured health outcomes, more research is needed into the correlates and determinants of our observed association. For example, our BMI trajectory results do not cast light on the importance of adiposity or lean muscle mass in relation to self-rated health at different ages.

In conclusion, BMI trajectories were associated with self-rated health, however the nature of the association depended on age. A higher level of BMI characterised those experiencing poor health among middle-aged participants, while among their older peers, poor health was associated with a steeper decline in BMI. On the other hand, ten years before reporting of poor health, cases had higher BMI across the entire age-range under examination. These results could not have been captured by studies using only one BMI measurement.

## Supporting information

## Acknowledgements

ELSA was developed by researchers from the University College London, the Institute of Fiscal Studies and the National Centre for Social Research.

## Contributors

AH, DBI, ASDL, and CCD conceived the study. AH and CCD wrote the first and successive drafts of the manuscript with substantial contribution from DBI and ASDL. AH carried out the statistical analysis. DBI and ASDL contributed to the data analysis. All authors revised the manuscript for important intellectual content. AH had full access to the data and take responsibility for the integrity of the data and the accuracy of the data analysis. AH is the guarantor.

## Funding

AH is supported by the Danish Diabetes Academy. The Danish Diabetes Academy is funded by the Novo Nordisk Foundation.

The funding for ELSA is provided by the National Institute of Aging in the USA, and a consortium of UK government departments coordinated by the Economic and Social Research Council. The funders of the English Longitudinal Study of Ageing and the UK Data Service do not bear any responsibility for the analyses or interpretations presented here.

## Competing interests

None declared.

## Patient consent

Not required.

## Ethics approval

ELSA was approved by the London Multicentre Research Ethics Committee (MREC/01/2/91),

## Data sharing statement

ELSA data are available for free upon registration to the UK Data Service (https://www.ukdataservice.ac.uk)

## Notes

#### Summary of Updates

New subgroup analyses, investigating the role of CVD and cancer, have been conducted and added to the manuscript. A new figure (Figure 2A) has been added to highlight differences in BMI 10 years before first report of poor health.

## REFERENCES

1. Bombak AE. Self-Rated Health and Public Health: A Critical Perspective. Front in Public Health 2013;1:15.

2. Bamia C, Orfanos P, Juerges H, et al. Self-rated health and all-cause and cause-specific mortality of older adults: Individual data meta-analysis of prospective cohort studies in the CHANCES Consortium. Maturitas 2017; 103:37–44.

3. Idler EL, Benyamini Y. Self-rated health and mortality: a review of twenty-seven community studies. J Health Soc Behav 1997;38:21–37.

4. Mavaddat N, Kinmonth AL, Sanderson S, et al. What determines Self-rated Health (SRH)? A cross-sectional study of SF-36 health domains in the EPIC-Norfolk cohort. J Epidemiol Community Health 2011;65:800–6.

5. Altman CE, Van Hook J, Hillemeier M. What Does Self-rated Health Mean? Changes and Variations in the Association of Obesity with Objective and Subjective Components Of Self-rated Health. J Health Social Behav 2016;57:39–58.

6. Stenholm S, Head J, Aalto V, et al. Body mass index as a predictor of healthy and disease-free life expectancy between ages 50 and 75: a multicohort study. Int J Obes (Lond) 2017;41:769–75.

7. Wang M, Yi Y, Roebothan B, et al. Body Mass Index Trajectories among Middle-Aged and Elderly Canadians and Associated Health Outcomes. J Environ Public Health 2016;2016:7014857.

8. Ruhunuhewa I, Adjibade M, Andreeva VA, et al. Prospective association between body mass index at midlife and healthy aging among French adults. Obesity (Silver Spring) 2017;25: 1254–62.

9. Giovannucci E. An Integrative Approach for Deciphering the Causal Associations of Physical Activity and Cancer Risk: The Role of Adiposity. J Natl Cancer Inst 2018;110:935–41.

10. Miller SL, Wolfe RR. The danger of weight loss in the elderly. J Nutr Health Aging 2008; 12:487–91.

11. Vanfleteren L, Lamprecht B, Wouters E, et al. The body mass index and chronic obstructive pulmonary disease in the BOLD study. Eur Respir J 2013;42(Suppl 57):P4652.

12. Steptoe A, Breeze E, Banks J, et al. Cohort profile: the English longitudinal study of ageing. Int J Epidemiol 2013;42:1640–8.

13. de Fine Olivarius N, Siersma VD, Køster-Rasmussen R, et al. Weight Changes following the Diagnosis of Type 2 Diabetes: The Impact of Recent and Past Weight History before Diagnosis. Results from the Danish Diabetes Care in General Practice (DCGP) Study. PloS One 2015;10:e0122219.

14. Chiolero A, Faeh D, Paccaud F, et al. Consequences of smoking for body weight, body fat distribution, and insulin resistance. Am J Clin Nutr 2008;87:801–9.

15. Wannamethee SG, Shaper AG, Walker M. Overweight and obesity and weight change in middle aged men: impact on cardiovascular disease and diabetes. J Epidemiology Community Health 2005;59:134–9.

16. Jackson SE, Williams K, Steptoe A, et al. The impact of a cancer diagnosis on weight change: findings from prospective, population-based cohorts in the UK and the US. BMC Cancer 2014;14:926.

17. Sirola J, Tuppurainen M, Rikkonen T, et al. Correlates and predictors of Self-rated health and ambulatory status among elderly women I Cross-sectional and 10 years population-based cohort study. Maturitas 2010; 65:244–52.

18. Svedberg P, Bardage C, Sandin S, et al. A prospective study of health, life-style and psychosocial predictors of Self-rated health. Eur J Epidemiol 2006;21:767–76.

19. Wagner DC, Short JL. Longitudinal predictors of Self-rated health and mortality in older adults. Prev Chronic Dis 2014;11:E93.

20. Dennerstein L, Dudley EC, Guthrie JR. Predictors of declining Self-rated health during the transition to menopause. J Psychosom Res 2003;54:147–53.

21. Simonsen MK, Hundrup YA, Gronbaek M, et al. A prospective study of the association between weight changes and Self-rated health. BMC Womens Health 2008;8:13.

22. Aune D, Sen A, Prasad M, et al. BMI and all cause mortality: systematic review and non-linear dose-response meta-analysis of 230 cohort studies with 3.74 million deaths among 30.3 million participants. BMJ 2016;353:i2156.

23. Feng X, Astell-Burt T. Impact of a type 2 diabetes diagnosis on mental health, quality of life, and social contacts: a longitudinal study. BMJ Open Diabetes Res Care 2017;5:e000198.

24. Shahar DR, Schultz R, Shahar A, et al. The effect of widowhood on weight change, dietary intake, and eating behavior in the elderly population. J Aging Health 2001;13:189–99.

