## Supplementary material for "Body mass index trajectories preceding first report of poor self-rated health: a longitudinal case-control analysis of the English Longitudinal Study of Ageing"

*11/12/2018*

### Introduction

This is an R Markdown file documenting the statistical analyses conducted for the manuscript in the title. Data used in the analyses is available from the UK Data Service (link).

### Session info

The reproducibility of the results might depend on the version of R and the used packages.

```
library(haven) # data import
library(carpenter) # table 1
library(magrittr) # utility functions
library(nlme) # mixed-effects model fitting
library(Epi) # confidence intervals for estimates and combined estimates
```

```
##
## Attaching package: 'Epi'

## The following object is masked from 'package:base':
##
##     merge.data.frame
```

```
library(kableExtra) # table formatting
```

```
## Warning: package 'kableExtra' was built under R version 3.4.4
```

```
sessionInfo()
```

```
## R version 3.4.3 (2017-11-30)
## Platform: x86_64-apple-darwin15.6.0 (64-bit)
## Running under: macOS High Sierra 10.13.6
##
## Matrix products: default
## BLAS: /Library/Frameworks/R.framework/Versions/3.4/Resources/lib/libRblas.0.dylib
## LAPACK: /Library/Frameworks/R.framework/Versions/3.4/Resources/lib/libRlapack.dylib
##
## locale:
## [1] en_GB.UTF-8/en_GB.UTF-8/en_GB.UTF-8/C/en_GB.UTF-8/en_GB.UTF-8
##
## attached base packages:
## [1] stats     graphics  grDevices utils     datasets  methods   base
##
## other attached packages:
## [1] kableExtra_0.9.0 Epi_2.12         nlme_3.1-131     magrittr_1.5
## [5] carpenter_0.2.1  haven_1.1.1
##
## loaded via a namespace (and not attached):
```

```
##  [1] Rcpp_0.12.16      pillar_1.1.0      compiler_3.4.3
##  [4] plyr_1.8.4        forcats_0.2.0     tools_3.4.3
##  [7] digest_0.6.14     viridisLite_0.2.0 evaluate_0.10.1
## [10] tibble_1.4.2      lattice_0.20-35   pkgconfig_2.0.1
## [13] rlang_0.1.6       Matrix_1.2-12     rstudioapi_0.7
## [16] cmprsk_2.2-7      yaml_2.1.16       parallel_3.4.3
## [19] stringr_1.2.0     httr_1.3.1        knitr_1.18
## [22] xml2_1.2.0        hms_0.4.1         rprojroot_1.3-2
## [25] grid_3.4.3        R6_2.2.2          survival_2.41-3
## [28] etm_0.6-2         rmarkdown_1.8     readr_1.1.1
## [31] backports_1.1.2   scales_0.5.0      htmltools_0.3.6
## [34] splines_3.4.3     MASS_7.3-47       rvest_0.3.2
## [37] colorspace_1.3-2  numDeriv_2016.8-1 stringi_1.1.6
## [40] munsell_0.4.3
```

### Data Import

Data files can be accessed after a short registration process at the UK Data Archive. Make sure to adjust the file paths, or that you have the data files in the same folder as the R script. The imported data are transformed into data.frames.

```
setwd("~/Research/ELSA2/UKDA-5050-stata/stata/stata13_se")

wave0_core <- read_dta("wave_0_common_variables_v2.dta")
wave0_core <- as.data.frame(wave0_core)

wave1_core <- read_dta("wave_1_core_data_v3.dta")
wave1_core <- as.data.frame(wave1_core)

wave2_core <- read_dta("wave_2_core_data_v4.dta")
wave2_core <- as.data.frame(wave2_core)

wave2_derived <- read_dta("wave_2_derived_variables.dta")
wave2_derived <- as.data.frame(wave2_derived)

wave2_nurse <- read_dta("wave_2_nurse_data_v2.dta")
wave2_nurse <- as.data.frame(wave2_nurse)

wave2_ces <- read_dta("wave_2_ifs_derived_variables.dta")
wave2_ces <- as.data.frame(wave2_ces)

wave3_core <- read_dta("wave_3_elsa_data_v4.dta")
wave3_core <- as.data.frame(wave3_core)

wave3_ces <- read_dta("wave_3_ifs_derived_variables.dta")
wave3_ces <- as.data.frame(wave3_ces)

wave4_core <- read_dta("wave_4_elsa_data_v3.dta")
wave4_core <- as.data.frame(wave4_core)

wave4_nurse <- read_dta("wave_4_nurse_data.dta")
wave4_nurse <- as.data.frame(wave4_nurse)
```

```r
wave4_ces <- read_dta("wave_4_ifs_derived_variables.dta")
wave4_ces <- as.data.frame(wave4_ces)

wave5_core <- read_dta("wave_5_elsa_data_v4.dta")
wave5_core <- as.data.frame(wave5_core)

wave5_ces <- read_dta("wave_5_ifs_derived_variables.dta")
wave5_ces <- as.data.frame(wave5_ces)

wave6_core <- read_dta("wave_6_elsa_data_v2.dta")
wave6_core <- as.data.frame(wave6_core)

wave6_nurse <- read_dta("wave_6_elsa_nurse_data_v2.dta")
wave6_nurse <- as.data.frame(wave6_nurse)

wave6_ces <- read_dta("wave_6_ifs_derived_variables.dta")
wave6_ces <- as.data.frame(wave6_ces)

wave7_core <- read_dta("wave_7_elsa_data.dta")
wave7_core <- as.data.frame(wave7_core)

wave7_ces <- read_dta("wave_7_ifs_derived_variables.dta")
wave7_ces <- as.data.frame(wave7_ces)
```

#### Select variables and merge waves

Prevalent CVD The new values represent the wave of diagnosis (0 means no diagnosis).

```r
hediaW1 <- c(paste0("hedia0",1:9), "hedia10")
hedimW1 <- paste0("hedim0",1:7)
hediaW2 <- paste0("hedia0",1:9)
hedimW2 <- paste0("hedim0",1:8)

# 3: heart attack
# 8: stroke
wave1_core$newCVD1 <- apply(wave1_core[,c(hediaW1,hedimW1)], 1, function(x) any(c(3,8) %in% x))
wave1_core$newCVD1 <- ifelse(wave1_core$newCVD1, 1, 99)

wave2_core$newCVD2 <- apply(wave2_core[,c(hediaW2,hedimW2)], 1, function(x) any(c(3,8) %in% x))
wave2_core$newCVD2 <- ifelse(wave2_core$newCVD2, 2, 99)

wave3_core$newCVD3 <- ifelse(wave3_core$hediami==1 | wave3_core$hediast==1, 3, 99)
table(wave3_core$newCVD3, useNA = "always")
```

```
##
##    3   99 <NA>
##  203 9568    0
```

```r
wave4_core$newCVD4 <- ifelse(wave4_core$hediami==1 | wave4_core$hediast==1, 4, 99)
table(wave4_core$newCVD4, useNA = "always")
```

```
##
##     4    99  <NA>
##   294 10756     0
```

```r
wave5_core$newCVD5 <- ifelse(wave5_core$hediami==1 | wave5_core$hediast==1, 5, 99)
table(wave5_core$newCVD5, useNA = "always")
```

```
##
##     5    99  <NA>
##   251 10023     0
```

```r
wave6_core$newCVD6 <- ifelse(wave6_core$hediami==1 | wave6_core$hediast==1, 6, 99)
table(wave6_core$newCVD6, useNA = "always")
```

```
##
##     6    99  <NA>
##   261 10340     0
```

```r
wave7_core$newCVD7 <- ifelse(wave7_core$hediami==1 | wave7_core$hediast==1, 7, 99)
table(wave7_core$newCVD7, useNA = "always")
```

```
##
##    7   99 <NA>
##  221 9445    0
```

Variables are selected from each wave and then renamed.

Time-invariant covariates (e.g. sex) are defined based on wave 1 as that is the official baseline wave of ELSA.

**Remark:** Self-rated health is a combination of two variables in wave 1, because the question was placed in two different positions in the questionnaire. However, each person was only asked once.

```r
wave0 <- wave0_core[, c("idauniq", "genhelf", "bmival", "ager")]
names(wave0) <- c("id", "srh0", "bmi0", "age0")

wave1_core$srh1 <- ifelse(wave1_core$hehelf < 0,
                          wave1_core$hehelfb,
                          wave1_core$hehelf)

wave1 <- wave1_core[, c("idauniq","srh1","indager","indsex","asoccls","hesmk","dimar","newCVD1")]
names(wave1) <- c("id", "srh1", "age1", "sex", "ses6", "smoke", "marital","newCVD1")


wave2_core$cancer <- ifelse(wave2_core$HeCana > 0, 1, 0)
wave2 <- wave2_core[, c("idauniq", "Hehelf", "indager", "HeACd", "cancer","newCVD2")]
names(wave2) <- c("id", "srh2", "age2", "dm2", "cancer2","newCVD2")

wave2_ces$cesd2 <- ifelse(wave2_ces$cesd_na==8, wave2_ces$cesd_sc, NA)
wave2_ces$id <- wave2_ces$idauniq
wave2_ces <- wave2_ces[,c("id", "cesd2")]

wave2_lung <- wave2_derived[, c("idauniq", "hediblu")]
names(wave2_lung) <- c("id", "lung2")

wave2_bmi <- wave2_nurse[, c("idauniq", "bmival")]
names(wave2_bmi) <- c("id", "bmi2")

wave3_core$cancer <- ifelse(wave3_core$hecanaa > 0, 1, 0)
wave3 <- wave3_core[, c("idauniq", "hegenh", "indager", "dheacd", "hediblu", "cancer","newCVD3")]
names(wave3) <- c("id", "srh3", "age3", "dm3", "lung3", "cancer3","newCVD3")
```

```r
wave3_ces$cesd3 <- ifelse(wave3_ces$cesd_na==8, wave3_ces$cesd_sc, NA)
wave3_ces$id <- wave3_ces$idauniq
wave3_ces <- wave3_ces[,c("id", "cesd3")]

wave4_core$cancer <- ifelse(wave4_core$hecanaa > 0, 1, 0)
wave4 <- wave4_core[, c("idauniq", "hehelf", "indager", "heacd", "hediblu", "cancer","newCVD4")]
names(wave4) <- c("id", "srh4", "age4", "dm4", "lung4", "cancer4","newCVD4")

wave4_bmi<- wave4_nurse[, c("idauniq", "bmival")]
names(wave4_bmi) <- c("id", "bmi4")

wave4_ces$cesd4 <- ifelse(wave4_ces$cesd_na==8, wave4_ces$cesd_sc, NA)
wave4_ces$id <- wave4_ces$idauniq
wave4_ces <- wave4_ces[,c("id", "cesd4")]

wave5_core$cancer <- ifelse(wave5_core$hecanaa > 0, 1, 0)
wave5 <- wave5_core[, c("idauniq", "hehelf", "indager", "heacd", "hediblu", "cancer","newCVD5")]
names(wave5) <- c("id", "srh5", "age5", "dm5", "lung5", "cancer5","newCVD5")

wave5_ces$cesd5 <- ifelse(wave5_ces$cesd_na==8, wave5_ces$cesd_sc, NA)
wave5_ces$id <- wave5_ces$idauniq
wave5_ces <- wave5_ces[,c("id", "cesd5")]

wave6_core$cancer <- ifelse(wave6_core$HeCanaa > 0, 1, 0)
wave6 <- wave6_core[, c("idauniq", "Hehelf", "indager", "HeACd", "hediblu", "cancer","newCVD6")]
names(wave6) <- c("id", "srh6", "age6", "dm6", "lung6", "cancer6","newCVD6")

wave6_bmi <- wave6_nurse[, c("idauniq", "BMIVAL")]
names(wave6_bmi) <- c("id", "bmi6")

wave6_ces$cesd6 <- ifelse(wave6_ces$cesd_na==8, wave6_ces$cesd_sc, NA)
wave6_ces$id <- wave6_ces$idauniq
wave6_ces <- wave6_ces[,c("id", "cesd6")]

wave7_core$cancer <- ifelse(wave7_core$HeCanaa > 0, 1, 0)
wave7 <- wave7_core[, c("idauniq", "Hehelf", "indager", "HeACd", "hediblu", "cancer","newCVD7")]
names(wave7) <- c("id", "srh7", "age7", "dm7", "lung7", "cancer7","newCVD7")

wave7_ces$cesd7 <- ifelse(wave7_ces$cesd_na==8, wave7_ces$cesd_sc, NA)
wave7_ces$id <- wave7_ces$idauniq
wave7_ces <- wave7_ces[,c("id", "cesd7")]

combined <- merge(wave0, wave1, by = "id", all = TRUE)
combined <- merge(combined, wave2, by = "id", all = TRUE)
combined <- merge(combined, wave3, by = "id", all = TRUE)
combined <- merge(combined, wave4, by = "id", all = TRUE)
combined <- merge(combined, wave5, by = "id", all = TRUE)
combined <- merge(combined, wave6, by = "id", all = TRUE)
combined <- merge(combined, wave7, by = "id", all = TRUE)

combined <- merge(combined, wave2_bmi, by = "id", all = TRUE)
combined <- merge(combined, wave4_bmi, by = "id", all = TRUE)
combined <- merge(combined, wave6_bmi, by = "id", all = TRUE)
```

```
combined <- merge(combined, wave2_lung, by = "id", all = TRUE)

combined <- merge(combined, wave2_ces, by = "id", all = TRUE)
combined <- merge(combined, wave3_ces, by = "id", all = TRUE)
combined <- merge(combined, wave4_ces, by = "id", all = TRUE)
combined <- merge(combined, wave5_ces, by = "id", all = TRUE)
combined <- merge(combined, wave6_ces, by = "id", all = TRUE)
combined <- merge(combined, wave7_ces, by = "id", all = TRUE)
```

### Clean variables

Missing values are coded with negative values in ELSA, these are transformed to NAs. Note, some NAs are generated (during merging data sets) for those who didn't participate in a specific wave. Therefore we saw a combination of negative values and NAs in the previous section. Now we change all of them to NA.

In waves 0 and 3, slightly different answers were given, therefore we merged categories.

Factor variables (e.g. sex: male/female) are defined.

```
srhVar <- paste0("srh", 0:7)
combined[, srhVar][combined[, srhVar] < 1] <- NA

combined$srh0[combined$srh0 == 4] <- 5
combined$srh3[combined$srh3 == 4] <- 5

bmiVar <- paste0("bmi", c(0, 2, 4, 6))
combined[, bmiVar][combined[, bmiVar] < 10] <- NA # using 10 here to exclude a value of 4

ageVar <- paste0("age", 0:7)
combined[, ageVar][combined[, ageVar] < 1] <- NA

combined[combined$sex < 1 | is.na(combined$sex), "sex"] <- NA
combined$sex <- factor(combined$sex, levels = c(1,2), labels = c("male","female"))

combined$marital[combined$marital < 1] <- NA
combined$neverMarried <- combined$marital == 1
combined$neverMarried <- factor(combined$neverMarried, levels = c(FALSE,TRUE),
                                labels = c("ever","never"))

combined$maritalStatus <- factor(combined$marital)
levels(combined$maritalStatus) <- c("single", "married", "married", "divorced", "divorced", "widowed")

combined$ses6[combined$ses6 < 1] <- NA
combined$ses6[combined$ses6 %in% c(7,8)] <- NA
combined$SES <- factor(combined$ses6)
levels(combined$SES) <- c("I&II","I&II","IIIN","IIIM","IV&V","IV&V")

combined$smoke[combined$smoke < 1] <- NA
combined$everSmoke <- factor(combined$smoke, levels = c(1,2), labels = c("yes","no"))

dmVar <- paste0("dm", 2:7)
combined$diabetes <- apply(combined[, dmVar], 1, function(x) 1 %in% x)
combined$diabetes <- factor(combined$diabetes, levels = c(FALSE,TRUE),
```

```
                            labels = c("no", "yes"))

lungVar <- paste0("lung", 2:7)
combined$lungAny <- apply(combined[, lungVar], 1, function(x) any(c(1,2) %in% x))
combined$lungAny <- factor(combined$lungAny, levels = c(FALSE,TRUE),
                            labels = c("no", "yes"))

cancerVar <- paste0("cancer", 2:7) # currently not in use
combined$cancer <- apply(combined[, cancerVar], 1, function(x) 1 %in% x)
combined$cancer <- factor(combined$cancer, levels = c(FALSE,TRUE),
                          labels = c("no", "yes"))

cvdVar <- paste0("newCVD",1:7)
combined$cvdWave <- apply(combined[,cvdVar], 1, min, na.rm=T) # warnings are generated for Inf
combined$cvdWave[combined$cvdWave==Inf] <- 99
```

#### Worst self-rated health up to a specific wave

We analyse first report of poor health, therefore we keep track of the worst reported health up to each wave. This variable only makes sense to use from wave 1. For those not having any valid measurements, the `max` function will generate `-Inf` values that we turn into NAs.

```
maxsrhVar <- paste0("maxsrh", 1:7)

combined$maxsrh1 <- combined$srh0

for(i in 2:7) {
  combined[[ maxsrhVar[i] ]] <- apply(combined[ , srhVar[1:i] ], 1,  max, na.rm=T)
}

combined[, maxsrhVar][combined[, maxsrhVar] == -Inf] <- NA
```

#### Core members

We consider only core members for this analysis.

```
coreIDs <- wave0_core[wave0_core$finstatw5 == 1, "idauniq"]
combined <- subset(combined, id %in% coreIDs)
nrow(combined)
```

```
## [1] 11205
```

#### Incident cases

conditions: srh == 5 in the current wave, but the previous maximum is < 5

maxsrhVar[i-1] is used, because there are only 7 of them (no maxsrh0) in other words srhVar[i] and maxsrhVar[i-1] refer to the same wave

```
first5Var <- paste0("first5_", 0:7)
combined$first5_0 <- combined$srh0 == 5

for(i in 2:8) {
```

```
  combined[[ first5Var[i] ]] <-
      combined[[ srhVar[i] ]] == 5 & combined[[ maxsrhVar[i-1] ]] < 5
}

matchTRUE <- function(x) match(TRUE, x) # function to find TRUE and return its index
combined$diagWave <- apply(combined[, first5Var[1:8]], 1, matchTRUE) - 1

combined$case0 <- !is.na(combined$diagWave)

combined$case <- combined$diagWave %in% 2:7

table(combined$case0,combined$case, useNA = "always")
```

```
##
##          FALSE TRUE <NA>
##   FALSE  8722    0    0
##   TRUE   1429 1054    0
##   <NA>      0    0    0
```

```
table(combined$case0, combined$diagWave, useNA = "always")
```

```
##
##            0    1    2    3    4    5    6    7 <NA>
##   FALSE    0    0    0    0    0    0    0    0 8722
##   TRUE  1012  417  302  186  171  161  132  102    0
##   <NA>     0    0    0    0    0    0    0    0    0
```

8722 never reported poor health.

1429 already reported at waves 0 and 1 (1012 & 417, respectively).

### Fix age variable

Above 90 years of age, 99 is used as value. We try to impute these based on information from the previous waves.

E.g. if someone has the following sequence 86, 88, 90, 99, then 92 is a good estimate instead of 99, assuming 2 year intervals between examinations. First we calculate the median age difference between consecutive waves. We will use that for calculating the next age when that is missing or coded as 99. Another scenario is when age is already 99 at the first wave. In this case, based on the life expectancy above 90 years, we substitute 99 with 92 years of age.

Note, that this method will impute the age of those who die during follow-up too.

```
diff10 <- as.numeric(median(combined$age1 - combined$age0, na.rm = TRUE))
diff21 <- as.numeric(median(combined$age2 - combined$age1, na.rm = TRUE))
diff32 <- as.numeric(median(combined$age3 - combined$age2, na.rm = TRUE))
diff43 <- as.numeric(median(combined$age4 - combined$age3, na.rm = TRUE))
diff54 <- as.numeric(median(combined$age5 - combined$age4, na.rm = TRUE))
diff65 <- as.numeric(median(combined$age6 - combined$age5, na.rm = TRUE))
diff76 <- as.numeric(median(combined$age7 - combined$age6, na.rm = TRUE))

combined[combined$age0==99, "age0"] <- 92

combined[combined$age1 %in% c(99,NA), "age1"] <-
    combined[combined$age1 %in% c(99,NA), "age0"] + diff10
```

```
combined[combined$age2 %in% c(99,NA), "age2"] <-
    combined[combined$age2 %in% c(99,NA), "age1"] + diff21

combined[combined$age3 %in% c(99,NA), "age3"] <-
    combined[combined$age3 %in% c(99,NA), "age2"] + diff32

combined[combined$age4 %in% c(99,NA), "age4"] <-
    combined[combined$age4 %in% c(99,NA), "age3"] + diff43

combined[combined$age5 %in% c(99,NA), "age5"] <-
    combined[combined$age5 %in% c(99,NA), "age4"] + diff54

combined[combined$age6 %in% c(99,NA), "age6"] <-
    combined[combined$age6 %in% c(99,NA), "age5"] + diff65

combined[combined$age7 %in% c(99,NA), "age7"] <-
    combined[combined$age7 %in% c(99,NA), "age6"] + diff76

# manual correction
# the row number (2195) is used not to disclose the id of the participant
combined[2195, paste0("age", 0:7)]
```

```
##      age0 age1 age2 age3 age4 age5 age6 age7
## 5680   57   60   62   64   81   68   70   73
```

```
combined[2195,"age4"] <- 66
```

Now, we are ready to calculate age at "diagnosis".

#### Age at first report of poor health

```
combined$diagAge <- NA
for(i in 1:nrow(combined)) {
    if (!is.na(combined$diagWave[i])) {
        if (combined$diagWave[i]==0) combined$diagAge[i] <- combined$age0[i]
        if (combined$diagWave[i]==1) combined$diagAge[i] <- combined$age1[i]
        if (combined$diagWave[i]==2) combined$diagAge[i] <- combined$age2[i]
        if (combined$diagWave[i]==3) combined$diagAge[i] <- combined$age3[i]
        if (combined$diagWave[i]==4) combined$diagAge[i] <- combined$age4[i]
        if (combined$diagWave[i]==5) combined$diagAge[i] <- combined$age5[i]
        if (combined$diagWave[i]==6) combined$diagAge[i] <- combined$age6[i]
        if (combined$diagWave[i]==7) combined$diagAge[i] <- combined$age7[i]
    }
}
```

### Time before first report of poor health

This is the time-scale in the model. Negative values represent measurements before the first report of poor health.

```r
tVar <- paste0("time",0:7)

for(i in 1:8) {
  combined[[ tVar[i] ]] <- combined[[ ageVar[i] ]] - combined[["diagAge"]]
}
```

#### Restriction based on number of BMI measurements

We create a long dataset, subset that and then extract the unique individuals to create the restricted wide data of cases.

```r
waves <- c(0,2,4,6)
timeBMI <- paste0("time",waves)

longVarCases <- c("id", "sex", "diagAge","diagWave", "SES", "neverMarried",
                  "maritalStatus","everSmoke", "diabetes", "lungAny","cancer","cvdWave",
                  timeBMI, bmiVar)

# keep only cases
wide0 <- subset(combined, case, select=longVarCases)
# reshape into long format
long0 <- reshape(wide0,
                 varying = list(timeBMI, bmiVar),
                 direction = "long",
                 idvar = "id",
                 sep = "",
                 v.names = c("t", "bmi"),
                 times = waves,
                 timevar = "wave")

# consider only measurements from the [-10, 0] period
long_t10_0 <- subset(long0, !is.na(bmi) & t<=0 & t>=-10)
length(unique(long_t10_0$id))
```

```
## [1] 983
```

```r
# include only those with >=2 measurements
tn <- table(long_t10_0$id)
tnID <- row.names(tn)
nMeasure_df <- data.frame(id = tnID, n = as.vector(tn))
long_t10_0 <- merge(long_t10_0, nMeasure_df)
long10min2 <- subset(long_t10_0, n > 1)
length(unique(long10min2$id))
```

```
## [1] 732
```

```r
nrow(long10min2)
```

```
## [1] 1681
```

```r
wideCases <- subset(combined, id %in% long10min2$id)
wideCases$diagSRH <- 5

cesdVar <- paste0("cesd",2:7)
for(i in seq_along(wideCases$id)) {
```

```
    wideCases$diagCESD[i] <- wideCases[i,cesdVar][ wideCases[i,"diagWave"]-1 ]
}

longCases <- long10min2
```

### Matching

We try to find 3 controls for each case within a +/-1 year age range and of the same sex among those who don't report poor health at time 0 (density sampling). A person cannot be a control twice for the same wave (meaning to contribute with the same times and BMI values), but can be a control again in another wave. Cases can also be sampled as controls in waves before their first report of poor self-rated health. See detailed comments when finding controls at wave 2.

When selecting potential controls, we make sure that they have at least two BMI measurements in the last 10 years by checking the previous two measurements (always within 10 years) and the one before only if that's within 10 years.

```
# initialize
set.seed(1234)
# number of controls per case
N <- 3
# maximum age difference during matching
age_thr <- 1

# wave 2

# select cases reporting poor health at wave 2
casesW2 <- subset(wideCases, diagWave==2, select = c("id", "sex", "age2"))
# vector to store already sampled control IDs to make sure everyone comes up max once
controlID_W2 <- c()
# loop through all the cases reporting poor health at wave 2
for (i in 1:nrow(casesW2)) {
    # store the sex and the age of the case
    sexI <- casesW2[i, "sex"]
    ageI <- casesW2[i, "age2"]
    # potential controls are identified
    potentialControl <- subset(combined, !(srh2 %in% c(NA,5)) &
                                   maxsrh2 < 5 &
                                   sex == sexI &
                                   abs(age2-ageI) <= age_thr &
                                   !(id %in% controlID_W2) &
                                   !is.na(bmi0) & !is.na(bmi2)
                              )
    # take a random sample of potential controls (N or if there are only less then
    # available then all of them)
    index <- sample(nrow(potentialControl),min(nrow(potentialControl), N))
    # keep only selected from potential controls
    control_df <- potentialControl[index,]
    # append the selected controls' ids to a vector
    controlID_W2 <- c(controlID_W2, control_df$id)
    # append the data of selected controls to the data frame storing all controls
    # in this wave
    if (i==1) {
```

```r
        controlsW2 <- control_df
    } else {
        controlsW2 <- rbind(controlsW2, control_df)
    }
}

# Number of cases
nrow(casesW2)
```

```
## [1] 187
```

```r
# Number of controls selected
nrow(controlsW2)
```

```
## [1] 561
```

```r
# wave 3
casesW3 <- subset(wideCases, diagWave==3, select = c("id", "sex", "age3"))
controlID_W3 <- c()
for (i in 1:nrow(casesW3)) {
    sexI <- casesW3[i, "sex"]
    ageI <- casesW3[i, "age3"]
    potentialControl <- subset(combined, !(srh3 %in% c(NA,5)) &
                                maxsrh3 < 5 &
                                sex == sexI &
                                abs(age3-ageI) <= age_thr &
                                !(id %in% controlID_W3) &
                                !is.na(bmi0) & !is.na(bmi2)
    )
    index <- sample(nrow(potentialControl), min(nrow(potentialControl), N))
    control_df <- potentialControl[index,]
    controlID_W3 <- c(controlID_W3, control_df$id)
    if (i==1) {
        controlsW3 <- control_df
    } else {
        controlsW3 <- rbind(controlsW3, control_df)
    }
}
# Number of cases
nrow(casesW3)
```

```
## [1] 125
```

```r
# Number of controls selected
nrow(controlsW3)
```

```
## [1] 374
```

```r
# wave 4
casesW4 <- subset(wideCases, diagWave==4, select = c("id", "sex", "age4"))
controlID_W4 <- c()
for (i in 1:nrow(casesW4)) {
    sexI <- casesW4[i, "sex"]
    ageI <- casesW4[i, "age4"]
    potentialControl <- subset(combined, !(srh4 %in% c(NA,5)) &
                                maxsrh4 < 5 &
                                sex == sexI &
```

```r
                                abs(age4-ageI) <= age_thr &
                                !(id %in% controlID_W4) &
                                (ifelse(age4-age0<=10, is.na(bmi0),TRUE) +
                                     is.na(bmi2) + is.na(bmi4)) <= 1
    )
    index <- sample(nrow(potentialControl), min(nrow(potentialControl), N))
    control_df <- potentialControl[index,]
    controlID_W4 <- c(controlID_W4, control_df$id)
    if (i==1) {
        controlsW4 <- control_df
    } else {
        controlsW4 <- rbind(controlsW4, control_df)
    }
}
# Number of cases
nrow(casesW4)
```

```
## [1] 132
```

```r
# Number of controls selected
nrow(controlsW4)
```

```
## [1] 396
```

```r
# wave 5
casesW5 <- subset(wideCases, diagWave==5, select = c("id", "sex", "age5"))
controlID_W5 <- c()
for (i in 1:nrow(casesW5)) {
    sexI <- casesW5[i, "sex"]
    ageI <- casesW5[i, "age5"]
    potentialControl <- subset(combined, !(srh5 %in% c(NA,5)) &
                                maxsrh5 < 5 &
                                sex == sexI &
                                abs(age5-ageI) <= age_thr &
                                !(id %in% controlID_W5) &
                                (ifelse(age5-age0<=10, is.na(bmi0), TRUE) +
                                     is.na(bmi2) + is.na(bmi4)) <= 1
    )
    index <- sample(nrow(potentialControl), min(nrow(potentialControl), N))
    control_df <- potentialControl[index,]
    controlID_W5 <- c(controlID_W5, control_df$id)
    if (i==1) {
        controlsW5 <- control_df
    } else {
        controlsW5 <- rbind(controlsW5, control_df)
    }
}
# Number of cases
nrow(casesW5)
```

```
## [1] 104
```

```r
# Number of controls selected
nrow(controlsW5)
```

```
## [1] 312
```

```r
# wave 6
casesW6 <- subset(wideCases, diagWave==6, select = c("id", "sex", "age6"))
controlID_W6 <- c()
for (i in 1:nrow(casesW6)) {
    sexI <- casesW6[i, "sex"]
    ageI <- casesW6[i, "age6"]
    potentialControl <- subset(combined, !(srh6 %in% c(NA,5)) &
                               maxsrh6 < 5 &
                               sex == sexI &
                               abs(age6-ageI) <= age_thr &
                               !(id %in% controlID_W6) &
                               (ifelse(age6-age2<=10, is.na(bmi2), TRUE) +
                                    is.na(bmi4) + is.na(bmi6)) <= 1
    )
    index <- sample(nrow(potentialControl),min(nrow(potentialControl),N))
    control_df <- potentialControl[index,]
    controlID_W6 <- c(controlID_W6, control_df$id)
    if (i==1) {
        controlsW6 <- control_df
    } else {
        controlsW6 <- rbind(controlsW6, control_df)
    }
}
# Number of cases
nrow(casesW6)
```

```
## [1] 101
```

```r
# Number of controls selected
nrow(controlsW6)
```

```
## [1] 303
```

```r
# wave 7
casesW7 <- subset(wideCases, diagWave==7, select = c("id", "sex", "age7"))
controlID_W7 <- c()
for (i in 1:nrow(casesW7)) {
    sexI <- casesW7[i, "sex"]
    ageI <- casesW7[i, "age7"]
    potentialControl <- subset(combined, !(srh7 %in% c(NA,5)) &
                               maxsrh7 < 5 &
                               sex == sexI &
                               abs(age7-ageI) <= age_thr &
                               !(id %in% controlID_W7) &
                               (ifelse(age7-age2<=10,is.na(bmi2),TRUE) +
                                    is.na(bmi4) + is.na(bmi6)) <= 1
    )
    index <- sample(nrow(potentialControl), min(nrow(potentialControl), N))
    control_df <- potentialControl[index,]
    controlID_W7 <- c(controlID_W7, control_df$id)
    if (i==1) {
        controlsW7 <- control_df
    } else {
        controlsW7 <- rbind(controlsW7, control_df)
    }
```

```
}
# Number of cases
nrow(casesW7)
```

```
## [1] 83
```

```
# Number of controls selected
nrow(controlsW7)
```

```
## [1] 249
```

There is one person in wave 3 who has only 2 controls.

#### Controls to long format

Before reshaping the data, we have to create `diagAge`, `diagSRH` and `diagWave` variables (age, self-rated health and wave at "diagnosis"). These come from the wave where the matching happens (the corresponding case's "diagnosis").

IDs are modified as controls can come up several times, but not within the same wave, so the wave number is appended like "123456_4" if person 123456 is a control in wave 4.

```
longVar <- c("id", "sex", "diagAge","diagWave","diagSRH", "SES", "neverMarried",
             "maritalStatus", "diabetes", "lungAny", "cancer", "everSmoke", "diagCESD","cvdWave",timeBMI
# wave 2
for(i in 1:8) {
    controlsW2[[ tVar[i] ]] <- controlsW2[[ ageVar[i] ]] - controlsW2[["age2"]]
}
controlsW2$diagAge <- controlsW2[["age2"]]
controlsW2$diagSRH <- controlsW2[["srh2"]]
controlsW2$diagCESD <- controlsW2[["cesd2"]] # new
controlsW2$diagWave <- 2
controlsW2 <- subset(controlsW2, select=longVar)
controlsW2$case <- FALSE
controlsW2$id <- paste0(controlsW2$id, "_2")
longControlsW2_0 <- reshape(controlsW2,
                            varying = list(timeBMI, bmiVar),
                            direction = "long",
                            idvar = "id",
                            sep = "",
                            v.names = c("t", "bmi"),
                            times = waves,
                            timevar = "wave")

longControlsW2 <- subset(longControlsW2_0, !is.na(bmi) & t<=0 & t>=-10)

# wave 3
for(i in 1:8) {
    controlsW3[[ tVar[i] ]] <- controlsW3[[ ageVar[i] ]] - controlsW3[["age3"]]
}
controlsW3$diagAge <- controlsW3[["age3"]]
controlsW3$diagSRH <- controlsW3[["srh3"]]
controlsW3$diagCESD <- controlsW3[["cesd3"]] # new
controlsW3$diagWave <- 3
controlsW3 <- subset(controlsW3, select=longVar)
```

```r
controlsW3$case <- FALSE
controlsW3$id <- paste0(controlsW3$id, "_3")
longControlsW3_0 <- reshape(controlsW3,
                            varying = list(timeBMI, bmiVar),
                            direction = "long",
                            idvar = "id",
                            sep = "",
                            v.names = c("t", "bmi"),
                            times = waves,
                            timevar = "wave")

longControlsW3 <- subset(longControlsW3_0, !is.na(bmi) & t<=0 & t>=-10)

# wave 4
for(i in 1:8) {
    controlsW4[[ tVar[i] ]] <- controlsW4[[ ageVar[i] ]] - controlsW4[["age4"]]
}
controlsW4$diagAge <- controlsW4[["age4"]]
controlsW4$diagSRH <- controlsW4[["srh4"]]
controlsW4$diagCESD <- controlsW4[["cesd4"]] # new
controlsW4$diagWave <- 4
controlsW4 <- subset(controlsW4, select=longVar)
controlsW4$case <- FALSE
controlsW4$id <- paste0(controlsW4$id, "_4")
longControlsW4_0 <- reshape(controlsW4,
                            varying = list(timeBMI, bmiVar),
                            direction = "long",
                            idvar = "id",
                            sep = "",
                            v.names = c("t", "bmi"),
                            times = waves,
                            timevar = "wave")

longControlsW4 <- subset(longControlsW4_0, !is.na(bmi) & t<=0 & t>=-10)

# wave 5
for(i in 1:8) {
    controlsW5[[ tVar[i] ]] <- controlsW5[[ ageVar[i] ]] - controlsW5[["age5"]]
}
controlsW5$diagAge <- controlsW5[["age5"]]
controlsW5$diagSRH <- controlsW5[["srh5"]]
controlsW5$diagCESD <- controlsW5[["cesd5"]] # new
controlsW5$diagWave <- 5
controlsW5 <- subset(controlsW5, select=longVar)
controlsW5$case <- FALSE
controlsW5$id <- paste0(controlsW5$id, "_5")
longControlsW5_0 <- reshape(controlsW5,
                            varying = list(timeBMI, bmiVar),
                            direction = "long",
                            idvar = "id",
                            sep = "",
                            v.names = c("t", "bmi"),
                            times = waves,
```

```
                            timevar = "wave")

longControlsW5 <- subset(longControlsW5_0, !is.na(bmi) & t<=0 & t>=-10)

# wave 6
for(i in 1:8) {
    controlsW6[[ tVar[i] ]] <- controlsW6[[ ageVar[i] ]]-controlsW6[["age6"]]
}
controlsW6$diagAge <- controlsW6[["age6"]]
controlsW6$diagSRH <- controlsW6[["srh6"]]
controlsW6$diagCESD <- controlsW6[["cesd6"]] # new
controlsW6$diagWave <- 6
controlsW6 <- subset(controlsW6, select=longVar)
controlsW6$case <- FALSE
controlsW6$id <- paste0(controlsW6$id, "_6")
longControlsW6_0 <- reshape(controlsW6,
                            varying = list(timeBMI, bmiVar),
                            direction = "long",
                            idvar = "id",
                            sep = "",
                            v.names = c("t", "bmi"),
                            times = waves,
                            timevar = "wave")

longControlsW6 <- subset(longControlsW6_0, !is.na(bmi) & t<=0 & t>=-10)

# wave 7
for(i in 1:8) {
    controlsW7[[ tVar[i] ]] <- controlsW7[[ ageVar[i] ]]-controlsW7[["age7"]]
}
controlsW7$diagAge <- controlsW7[["age7"]]
controlsW7$diagSRH <- controlsW7[["srh7"]]
controlsW7$diagCESD <- controlsW7[["cesd7"]] # new
controlsW7$diagWave <- 7
controlsW7 <- subset(controlsW7, select=longVar)
controlsW7$case <- FALSE
controlsW7$id <- paste0(controlsW7$id, "_7")
longControlsW7_0 <- reshape(controlsW7,
                            varying = list(timeBMI, bmiVar),
                            direction = "long",
                            idvar = "id",
                            sep = "",
                            v.names = c("t", "bmi"),
                            times = waves,
                            timevar = "wave")

longControlsW7 <- subset(longControlsW7_0, !is.na(bmi) & t<=0 & t>=-10)
```

### Merging data sets

Data sets are merged and variables are harmonized.

```r
# merge controls
wideControls <- rbind(
    controlsW2,
    controlsW3,
    controlsW4,
    controlsW5,
    controlsW6,
    controlsW7
)

longControls <- rbind(
    longControlsW2,
    longControlsW3,
    longControlsW4,
    longControlsW5,
    longControlsW6,
    longControlsW7
)

# wide
wideFinal <- rbind(
    wideControls,
    wideCases[, names(wideControls)]
)

wideFinal$case <- factor(wideFinal$case, levels = c(FALSE, TRUE),
                         labels = c("control", "case"))
wideFinal$diagAge <- as.numeric(wideFinal$diagAge)
wideFinal$diagCESD <- as.numeric(wideFinal$diagCESD)
wideFinal$diagSRH <- factor(wideFinal$diagSRH)

wideFinal$diagCVD <- wideFinal$cvdWave <= wideFinal$diagWave


wideFinal$diagCVD <- factor(wideFinal$diagCVD, levels = c(FALSE, TRUE),
                         labels = c("no", "yes"))


# long
longCases$case <- TRUE

commonLongVar <- intersect(names(longControls), names(longCases))

longFinal <- rbind(
    longControls[, commonLongVar],
    longCases[, commonLongVar]
)

longFinal$case <- factor(longFinal$case, levels = c(FALSE, TRUE),
                         labels = c("control", "case"))

longFinal$diagAge <- as.numeric(longFinal$diagAge)
```

```
longFinal$diagCVD <- longFinal$id %in% wideFinal[wideFinal$diagCVD,"id"]
longFinal$diagCVD <- factor(longFinal$diagCVD, levels = c(FALSE, TRUE),
                         labels = c("no", "yes"))
```

#### Diabetes, cancer and chronic lung disease at time=0

These variables consider questionnaire-based information on diabetes and chronic lung disease status up to the wave of time=0. If someone ever reported e.g. diabetes before time=0, he/she is considered to have it even if it's not reported in later waves.

```
# wideFinal has new IDs for controls
wideFinal$id_orig <- substr(wideFinal$id, 1, 6)

# diabetes
diabData <- merge(wideFinal[, c("id","id_orig", "diabetes", "diagWave")],
                  combined[, c("id",paste0("dm",2:7))], all.x=T, all.y=F,by.x="id_orig",by.y="id")

diabData$diagDM <- NA
for(i in seq_along(diabData$id)) {
    diabData$diagDM[i] <- 1 %in% diabData[i, 5:(diabData[i,"diagWave"]+3)]
}

diabAtDiagnosis_ID <- diabData[diabData$diagDM,"id"]

# some might have it in wave 1
hedia <- paste0("hedia0",1:7)
baselineDM_id <- wave1_core[apply(wave1_core[,hedia], 1, function(x) 7 %in% x), "idauniq"]

wideFinal$diagDM <- wideFinal$id %in% union(diabAtDiagnosis_ID, baselineDM_id)
longFinal$diagDM <- longFinal$id %in% union(diabAtDiagnosis_ID, baselineDM_id)

wideFinal$diagDM <- factor(wideFinal$diagDM, levels = c(FALSE,TRUE),
                            labels = c("no", "yes"))
longFinal$diagDM <- factor(longFinal$diagDM, levels = c(FALSE,TRUE),
                            labels = c("no", "yes"))
# cancer
cancerData <- merge(wideFinal[, c("id","id_orig", "cancer", "diagWave")],
                  combined[, c("id",paste0("cancer",2:7))], all.x=T, all.y=F,by.x="id_orig",by.y="id")

cancerData$diagCancer <- NA
for(i in seq_along(cancerData$id)) {
    cancerData$diagCancer[i] <- 1 %in% cancerData[i, 5:(cancerData[i,"diagWave"]+3)]
}

cancerAtDiagnosis_ID <- cancerData[cancerData$diagCancer,"id"]

# some might have it in wave 1
baselineCancer_id <- wave1_core[wave1_core$hecana>0, "idauniq"]
wideFinal$diagCancer <- wideFinal$id %in% union(cancerAtDiagnosis_ID, baselineCancer_id)
longFinal$diagCancer <- longFinal$id %in% union(cancerAtDiagnosis_ID, baselineCancer_id)

wideFinal$diagCancer <- factor(wideFinal$diagCancer, levels = c(FALSE,TRUE),
                            labels = c("no", "yes"))
```

```
longFinal$diagCancer <- factor(longFinal$diagCancer, levels = c(FALSE,TRUE),
                              labels = c("no", "yes"))

# lung disease
lungData <- merge(wideFinal[, c("id","id_orig", "lungAny", "diagWave")],
                  combined[, c("id",paste0("lung",2:7))], all.x=T, all.y=F,by.x="id_orig",by.y="id")

lungData$diagLung <- NA
for(i in seq_along(lungData$id)) {
    lungData$diagLung[i] <- 1 %in% lungData[i, 5:(lungData[i,"diagWave"]+3)]
}

lungAtDiagnosis_ID <- lungData[lungData$diagLung,"id"]

wideFinal$diagLung <- wideFinal$id %in% lungAtDiagnosis_ID
longFinal$diagLung <- longFinal$id %in% lungAtDiagnosis_ID

wideFinal$diagLung <- factor(wideFinal$diagLung, levels = c(FALSE,TRUE),
                              labels = c("no", "yes"))
longFinal$diagLung <- factor(longFinal$diagLung, levels = c(FALSE,TRUE),
                              labels = c("no", "yes"))
```

### Table 1

```
wideFinal  %>%
    outline_table('case') %>%
    add_rows('diagAge', stat_medianIQR) %>%
    add_rows('diagCESD', stat_medianIQR) %>%
    add_rows('sex', stat_nPct) %>%
    add_rows('SES', stat_nPct) %>%
    add_rows('maritalStatus', stat_nPct) %>%
    add_rows('everSmoke', stat_nPct) %>%
    add_rows('diagDM', stat_nPct) %>%
    add_rows('diagCVD', stat_nPct) %>%
    add_rows('diagSRH', stat_nPct) %>%
    add_rows('diagLung', stat_nPct) %>%
    add_rows('diagCancer', stat_nPct) %>%
    build_table()
```

```
## Warning: attributes are not identical across measure variables;
## they will be dropped
```

| Variables | control | case |
|---|---|---|
| diagAge | 72.0 (64.0-80.0) | 73.0 (64.0-80.0) |
| diagCESD | 1.0 (0.0-2.0) | 3.0 (1.0-5.0) |
| diagCancer | | |
| - no | 2073 (94.4%) | 579 (79.1%) |
| - yes | 122 (5.6%) | 153 (20.9%) |
| diagCVD | | |
| - no | 1957 (89.2%) | 575 (78.6%) |
| - yes | 238 (10.8%) | 157 (21.4%) |
| diagDM | | |

| Variables | control | case |
| --- | --- | --- |
| - no | 1996 (90.9%) | 590 (80.6%) |
| - yes | 199 (9.1%) | 142 (19.4%) |
| diagLung | | |
| - no | 2081 (94.8%) | 621 (84.8%) |
| - yes | 114 (5.2%) | 111 (15.2%) |
| diagSRH | | |
| - 1 | 326 (14.9%) | |
| - 2 | 745 (33.9%) | |
| - 3 | 758 (34.5%) | |
| - 4 | 366 (16.7%) | |
| - 5 | | 732 (100%) |
| everSmoke | | |
| - no | 832 (38%) | 214 (29.3%) |
| - yes | 1355 (62%) | 517 (70.7%) |
| maritalStatus | | |
| - divorced | 201 (9.2%) | 99 (13.5%) |
| - married | 1461 (66.6%) | 443 (60.5%) |
| - single | 113 (5.1%) | 39 (5.3%) |
| - widowed | 420 (19.1%) | 151 (20.6%) |
| SES | | |
| - I&II | 812 (37.7%) | 218 (30.7%) |
| - IIIM | 366 (17%) | 131 (18.5%) |
| - IIIN | 569 (26.4%) | 157 (22.1%) |
| - IV&V | 405 (18.8%) | 203 (28.6%) |
| sex | | |
| - female | 1229 (56%) | 410 (56%) |
| - male | 966 (44%) | 322 (44%) |

```r
# n
table(wideFinal$case)
```

```
##
## control    case
##    2195     732
```

```r
# average number of measurements
table(longFinal$case)/table(wideFinal$case)
```

```
##
##  control     case
## 2.356720 2.296448
```

```r
# distribution of measurement numbers
with(subset(longFinal, case=="control"), prop.table(table(table(id))))
```

```
##
##            2            3            4
## 0.6437357631 0.3558086560 0.0004555809
```

```r
with(subset(longFinal, case=="case"), prop.table(table(table(id))))
```

```
##
##         2         3
## 0.7035519 0.2964481
```

```
# summary of first measurement times
sortTimeData <- longFinal[order(longFinal$t), c("id","t", "case")]
summary(sortTimeData[match(unique(sortTimeData$id),sortTimeData$id),"t"])
```

```
##    Min. 1st Qu.  Median    Mean 3rd Qu.    Max.
## -10.000  -9.000  -7.000  -7.044  -6.000  -2.000
```

### Modeling decisions

First, we investigated the effects of age at diagnosis and sex on BMI level and slope.

```
model0 <- lme(bmi ~ t*sex + t*diagAge,
              data = longFinal,
              random=~t|id,
              na.action = na.omit,
              method="ML",
              control = lmeControl(msMaxIter = 250, opt="optim"))

kable(ci.lin(model0)[, c(1,5,6,4)], "latex", booktabs = T) %>%
    kable_styling(latex_options = c("striped", "hold_position", "repeat_header"))
```

| | Estimate | 2.5% | 97.5% | P |
|---|---|---|---|---|
| (Intercept) | 32.1408640 | 30.7573523 | 33.5243757 | 0.0000000 |
| t | 0.5318226 | 0.4313606 | 0.6322846 | 0.0000000 |
| sexfemale | 0.0160590 | -0.3506470 | 0.3827650 | 0.9316000 |
| diagAge | -0.0586062 | -0.0773350 | -0.0398774 | 0.0000000 |
| t:sexfemale | 0.0029913 | -0.0229574 | 0.0289401 | 0.8212472 |
| t:diagAge | -0.0069772 | -0.0083270 | -0.0056273 | 0.0000000 |

Only age had an effect on the trajectories (both on level and slope), therefore we investigate the role of age, but not sex further.

### Code to create figures

Figures are created as functions so that they can be used for both exporting and presenting in the PDF.

```
x <- seq(-9, 0, 0.5)
ageAtDiag <- seq(60, 90, 1)
colIndex <- c(1,5,6)

modelFinal <- lme(bmi ~ t*diagAge*case,
                  data = longFinal,
                  random=~t|id,
                  na.action = na.omit,
                  method="ML",
                  control = lmeControl(msMaxIter = 250, opt="optim"))

# level
levelPlot <- function() {
    par(mar=c(4.5,4.5,1.1,1.1))
    plot(c(-9),c(-9),type="l",xlab="Age at first report of poor health (year)",
     ylab=expression(paste('BMI (kg/m'^'2'*')')),xlim=c(60,90),ylim=c(25,31),
```

```
     lwd=3,col="red",bty="n")
    text(88, 30.5, "B")
    # non-case
    matlines(ageAtDiag, ci.lin(modelFinal, ctr.mat = cbind(1, 0, ageAtDiag, 0, 0, 0, 0, 0))[,colIndex],
             col="blue", lwd = c(2,1,1), lty = c(1,3,3))
    # cases
    matlines(ageAtDiag, ci.lin(modelFinal, ctr.mat = cbind(1, 0, ageAtDiag, 1, 0, 0, ageAtDiag, 0))[,co
             col = "red", lwd = c(2,1,1), lty = c(1,3,3))
}

levelPlot10y <- function() {
    par(mar=c(4.5,4.5,1.1,1.1))
    plot(c(-9),c(-9),type="l",xlab="Age at first report of poor health (year)",
     ylab=expression(paste('BMI (kg/m'^'2'*')')),xlim=c(60,90),ylim=c(25,31),
     lwd=3,col="red",bty="n")
    text(88, 30.5, "A")
    # non-case
    matlines(ageAtDiag,
             ci.lin(modelFinal,
                                  ctr.mat = cbind(1, -10, ageAtDiag, 0, -10*ageAtDiag, 0, 0, 0))[,colIndex
             col="blue", lwd = c(2,1,1), lty = c(1,3,3))
    # cases
    matlines(ageAtDiag,
             ci.lin(modelFinal,
                                  ctr.mat = cbind(1, -10, ageAtDiag, 1, -10*ageAtDiag, -10, ageAtDiag, -10
             col = "red", lwd = c(2,1,1), lty = c(1,3,3))
}

# slopes
slopePlot <- function() {
    par(mar=c(4.5,4.5,1.1,1.1))
    plot(c(-9),c(-9),type="l",xlab="Age at first report of poor health (year)",
     ylab=expression(paste('BMI change (kg/m'^'2'*' per decade)')),xlim=c(60,90),ylim=c(-2.5,2),
     lwd=3,col="red",bty="n",yaxt="n")
    lines(c(50,90), c(0,0), col="gray")
    axis(2,at=seq(-2.5,2,0.5),las=1)
    text(88, 1.625, "C")
    # non-case
    matlines(ageAtDiag, 10*ci.lin(modelFinal, ctr.mat = cbind(0, 1, 0, 0, ageAtDiag, 0, 0, 0))[,colInde
             col="blue", lwd = c(2,1,1), lty = c(1,3,3))
    # cases
    matlines(ageAtDiag, 10*ci.lin(modelFinal, ctr.mat = cbind(0, 1, 0, 0, ageAtDiag, 1, 0, ageAtDiag))[
             col = "red", lwd = c(2,1,1), lty = c(1,3,3))
}

mainPlot <- function() {
    par(mar=c(4.5,4.5,1.1,1.1))
    plot(c(-9),c(-9),type="l",xlab="Age (year)",
     ylab=expression(paste('BMI (kg/m'^'2'*')')),xlim=c(50,90),ylim=c(25,31),
     lwd=3,col="red",bty="n")
    text(88, 30.5, "A")
    for(endAge in seq(60,90,10)) {
        controlMatrix <- cbind(1, x, endAge, 0, endAge*x, 0, 0, 0)
```

```
        controlPoint <- cbind(1, 0, endAge, 0, 0, 0, 0, 0)
        caseMatrix <- cbind(1, x, endAge, 1, endAge*x, x, endAge, endAge*x)
        casePoint <- cbind(1, 0, endAge, 1, 0, 0, endAge, 0)
        # non-case
        matlines(endAge + x, ci.lin(modelFinal, ctr.mat = controlMatrix)[,colIndex],
                 col="blue", lwd = c(2,1,1), lty = c(1,3,3))
        points(endAge, ci.lin(modelFinal, ctr.mat = controlPoint)[,1],
               pch=1, col="blue")

        # cases
        matlines(endAge + x, ci.lin(modelFinal, ctr.mat = caseMatrix)[,colIndex],
                 col = "red", lwd = c(2,1,1), lty = c(1,3,3))
        points(endAge, ci.lin(modelFinal, ctr.mat = casePoint)[,1],
               pch=16, col = "red")
    }
}

# SES-adjusted
modelSES <- lme(bmi ~ t*diagAge*case + SES,
                   data = longFinal,
                   random=~t|id,
                   na.action = na.omit,
                   method="ML",
                   control = lmeControl(msMaxIter = 250, opt="optim"))

sesPlot <- function() {
    par(mar=c(4.5,4.5,1.1,1.1))
    plot(c(-9),c(-9),type="l",xlab="Age (year)",
     ylab=expression(paste('BMI (kg/m'^'2'*')')),xlim=c(50,90),ylim=c(25,31),
     lwd=3,col="red",bty="n")
    for(endAge in seq(60,90,10)) {
        controlMatrix <- cbind(1,x,endAge, 0,0,0,0, endAge*x, 0,0, 0)
        controlPoint <- cbind(1,0,endAge, 0, 0,0,0, 0, 0,0, 0)
        caseMatrix <- cbind(1,x,endAge, 1,0,0,0, endAge*x, x, endAge,x*endAge)
        casePoint <- cbind(1,0,endAge, 1,0,0, 0, 0,0, endAge,0)
        # non-case
        matlines(endAge + x, ci.lin(modelSES, ctr.mat = controlMatrix)[,colIndex],
                 col="blue", lwd = c(2,1,1), lty = c(1,3,3))
        points(endAge, ci.lin(modelSES, ctr.mat = controlPoint)[,1],
               pch=1, col="blue")

        # cases
        matlines(endAge + x, ci.lin(modelSES, ctr.mat = caseMatrix)[,colIndex],
                 col = "red", lwd = c(2,1,1), lty = c(1,3,3))
        points(endAge, ci.lin(modelSES, ctr.mat = casePoint)[,1],
               pch=16, col = "red")
    }
}

# no diabetes
modelDM <- lme(bmi ~ t*diagAge*case,
                   data = subset(longFinal, diagDM=="no"),
                   random=~t|id,
```

```
                   na.action = na.omit,
                   method="ML",
                   control = lmeControl(msMaxIter = 250, opt="optim"))


dmPlot <- function() {
    par(mar=c(4.5,4.5,1.1,1.1))
    plot(c(-9),c(-9),type="l",xlab="Age (year)",
     ylab=expression(paste('BMI (kg/m'^'2'*')')),xlim=c(50,90),ylim=c(25,31),
     lwd=3,col="red",bty="n")
    text(88, 30.5, "C")
    for(endAge in seq(60,90,10)) {
        controlMatrix <- cbind(1, x, endAge, 0, endAge*x, 0, 0, 0)
        controlPoint <- cbind(1, 0, endAge, 0, 0, 0, 0, 0)
        caseMatrix <- cbind(1, x, endAge, 1, endAge*x, x, endAge, endAge*x)
        casePoint <- cbind(1, 0, endAge, 1, 0, 0, endAge, 0)
        # non-case
        matlines(endAge + x, ci.lin(modelDM, ctr.mat = controlMatrix)[,colIndex],
                 col="blue", lwd = c(2,1,1), lty = c(1,3,3))
        points(endAge, ci.lin(modelDM, ctr.mat = controlPoint)[,1],
               pch=1, col="blue")

        # cases
        matlines(endAge + x, ci.lin(modelDM, ctr.mat = caseMatrix)[,colIndex],
                 col = "red", lwd = c(2,1,1), lty = c(1,3,3))
        points(endAge, ci.lin(modelDM, ctr.mat = casePoint)[,1],
               pch=16, col = "red")
    }
}

# no lung disease
modelLung <- lme(bmi ~ t*diagAge*case,
                   data = subset(longFinal, diagLung=="no"),
                   random=~t|id,
                   na.action = na.omit,
                   method="ML",
                   control = lmeControl(msMaxIter = 250, opt="optim"))


lungPlot <- function() {
    par(mar=c(4.5,4.5,1.1,1.1))
    plot(c(-9),c(-9),type="l",xlab="Age (year)",
     ylab=expression(paste('BMI (kg/m'^'2'*')')),xlim=c(50,90),ylim=c(25,31),
     lwd=3,col="red",bty="n")
    text(88, 30.5, "F")
    for(endAge in seq(60,90,10)) {
        controlMatrix <- cbind(1, x, endAge, 0, endAge*x, 0, 0, 0)
        controlPoint <- cbind(1, 0, endAge, 0, 0, 0, 0, 0)
        caseMatrix <- cbind(1, x, endAge, 1, endAge*x, x, endAge, endAge*x)
        casePoint <- cbind(1, 0, endAge, 1, 0, 0, endAge, 0)
        # non-case
        matlines(endAge + x, ci.lin(modelLung, ctr.mat = controlMatrix)[,colIndex],
                 col="blue", lwd = c(2,1,1), lty = c(1,3,3))
```

```r
        points(endAge, ci.lin(modelLung, ctr.mat = controlPoint)[,1],
               pch=1, col="blue")

        # cases
        matlines(endAge + x, ci.lin(modelLung, ctr.mat = caseMatrix)[,colIndex],
                 col = "red", lwd = c(2,1,1), lty = c(1,3,3))
        points(endAge, ci.lin(modelLung, ctr.mat = casePoint)[,1],
               pch=16, col = "red")
    }
}

# never smokers
model_neverSMK <- lme(bmi ~ t*diagAge*case,
                   data = subset(longFinal, everSmoke=="no"),
                   random=~t|id,
                   na.action = na.omit,
                   method="ML",
                   control = lmeControl(msMaxIter = 250, opt="optim"))

smokePlot <- function() {
    par(mar=c(4.5,4.5,1.1,1.1))
    plot(c(-9),c(-9),type="l",xlab="Age (year)",
     ylab=expression(paste('BMI (kg/m'^'2'*')')),xlim=c(50,90),ylim=c(25,31),
     lwd=3,col="red",bty="n")
    text(88, 30.5, "B")
    for(endAge in seq(60,90,10)) {
        controlMatrix <- cbind(1, x, endAge, 0, endAge*x, 0, 0, 0)
        controlPoint <- cbind(1, 0, endAge, 0, 0, 0, 0, 0)
        caseMatrix <- cbind(1, x, endAge, 1, endAge*x, x, endAge, endAge*x)
        casePoint <- cbind(1, 0, endAge, 1, 0, 0, endAge, 0)
        # non-case
        matlines(endAge + x, ci.lin(model_neverSMK, ctr.mat = controlMatrix)[,colIndex],
                 col="blue", lwd = c(2,1,1), lty = c(1,3,3))
        points(endAge, ci.lin(model_neverSMK, ctr.mat = controlPoint)[,1],
               pch=1, col="blue")

        # cases
        matlines(endAge + x, ci.lin(model_neverSMK, ctr.mat = caseMatrix)[,colIndex],
                 col = "red", lwd = c(2,1,1), lty = c(1,3,3))
        points(endAge, ci.lin(model_neverSMK, ctr.mat = casePoint)[,1],
               pch=16, col = "red")
    }
}

# no cancer at diag
model_Cancer <- lme(bmi ~ t*diagAge*case,
                   data = subset(longFinal, diagCancer=="no"),
                   random=~t|id,
                   na.action = na.omit,
                   method="ML",
                   control = lmeControl(msMaxIter = 250, opt="optim"))

cancerPlot <- function() {
```

```
    par(mar=c(4.5,4.5,1.1,1.1))
    plot(c(-9),c(-9),type="l",xlab="Age (year)",
     ylab=expression(paste('BMI (kg/m'^'2'*')')),xlim=c(50,90),ylim=c(25,31),
     lwd=3,col="red",bty="n")
    text(88, 30.5, "E")
    for(endAge in seq(60,90,10)) {
        controlMatrix <- cbind(1, x, endAge, 0, endAge*x, 0, 0, 0)
        controlPoint <- cbind(1, 0, endAge, 0, 0, 0, 0, 0)
        caseMatrix <- cbind(1, x, endAge, 1, endAge*x, x, endAge, endAge*x)
        casePoint <- cbind(1, 0, endAge, 1, 0, 0, endAge, 0)
        # non-case
        matlines(endAge + x, ci.lin(model_Cancer, ctr.mat = controlMatrix)[,colIndex],
                 col="blue", lwd = c(2,1,1), lty = c(1,3,3))
        points(endAge, ci.lin(model_Cancer, ctr.mat = controlPoint)[,1],
               pch=1, col="blue")

        # cases
        matlines(endAge + x, ci.lin(model_Cancer, ctr.mat = caseMatrix)[,colIndex],
                 col = "red", lwd = c(2,1,1), lty = c(1,3,3))
        points(endAge, ci.lin(model_Cancer, ctr.mat = casePoint)[,1],
               pch=16, col = "red")
    }
}

# no CVD at diag
model_CVD <- lme(bmi ~ t*diagAge*case,
                   data = subset(longFinal, diagCVD=="no"),
                   random=~t|id,
                   na.action = na.omit,
                   method="ML",
                   control = lmeControl(msMaxIter = 250, opt="optim"))

cvdPlot <- function() {
    par(mar=c(4.5,4.5,1.1,1.1))
    plot(c(-9),c(-9),type="l",xlab="Age (year)",
     ylab=expression(paste('BMI (kg/m'^'2'*')')),xlim=c(50,90),ylim=c(25,31),
     lwd=3,col="red",bty="n")
    text(88, 30.5, "D")
    for(endAge in seq(60,90,10)) {
        controlMatrix <- cbind(1, x, endAge, 0, endAge*x, 0, 0, 0)
        controlPoint <- cbind(1, 0, endAge, 0, 0, 0, 0, 0)
        caseMatrix <- cbind(1, x, endAge, 1, endAge*x, x, endAge, endAge*x)
        casePoint <- cbind(1, 0, endAge, 1, 0, 0, endAge, 0)
        # non-case
        matlines(endAge + x, ci.lin(model_CVD, ctr.mat = controlMatrix)[,colIndex],
                 col="blue", lwd = c(2,1,1), lty = c(1,3,3))
        points(endAge, ci.lin(model_CVD, ctr.mat = controlPoint)[,1],
               pch=1, col="blue")

        # cases
        matlines(endAge + x, ci.lin(model_CVD, ctr.mat = caseMatrix)[,colIndex],
                 col = "red", lwd = c(2,1,1), lty = c(1,3,3))
        points(endAge, ci.lin(model_CVD, ctr.mat = casePoint)[,1],
```

```
                pch=16, col = "red")
    }
}

# none of CVD, cancer, DM, lung at diagnosis
subsetNone <- subset(longFinal, diagCVD=="no" & diagLung=="no" & diagCancer == "no" & diagDM == "no")
subsetNone_wide <- subset(wideFinal, diagCVD=="no" & diagLung=="no" & diagCancer == "no" & diagDM == "no
model_none <- lme(bmi ~ t*diagAge*case,
                   data = subsetNone,
                   random=~t|id,
                   na.action = na.omit,
                   method="ML",
                   control = lmeControl(msMaxIter = 250, opt="optim"))

nonePlot <- function() {
    par(mar=c(4.5,4.5,1.1,1.1))
    plot(c(-9),c(-9),type="l",xlab="Age (year)",
     ylab=expression(paste('BMI (kg/m'^'2'*')')),xlim=c(50,90),ylim=c(25,31),
     lwd=3,col="red",bty="n")
    text(88, 30.5, "")
    for(endAge in seq(60,90,10)) {
        controlMatrix <- cbind(1, x, endAge, 0, endAge*x, 0, 0, 0)
        controlPoint <- cbind(1, 0, endAge, 0, 0, 0, 0, 0)
        caseMatrix <- cbind(1, x, endAge, 1, endAge*x, x, endAge, endAge*x)
        casePoint <- cbind(1, 0, endAge, 1, 0, 0, endAge, 0)
        # non-case
        matlines(endAge + x, ci.lin(model_none, ctr.mat = controlMatrix)[,colIndex],
                 col="blue", lwd = c(2,1,1), lty = c(1,3,3))
        points(endAge, ci.lin(model_none, ctr.mat = controlPoint)[,1],
               pch=1, col="blue")

        # cases
        matlines(endAge + x, ci.lin(model_none, ctr.mat = caseMatrix)[,colIndex],
                 col = "red", lwd = c(2,1,1), lty = c(1,3,3))
        points(endAge, ci.lin(model_none, ctr.mat = casePoint)[,1],
               pch=16, col = "red")
    }
}
```

### Figure 2

```
par(mfrow=c(1,3))
levelPlot10y()
levelPlot()
slopePlot()
```

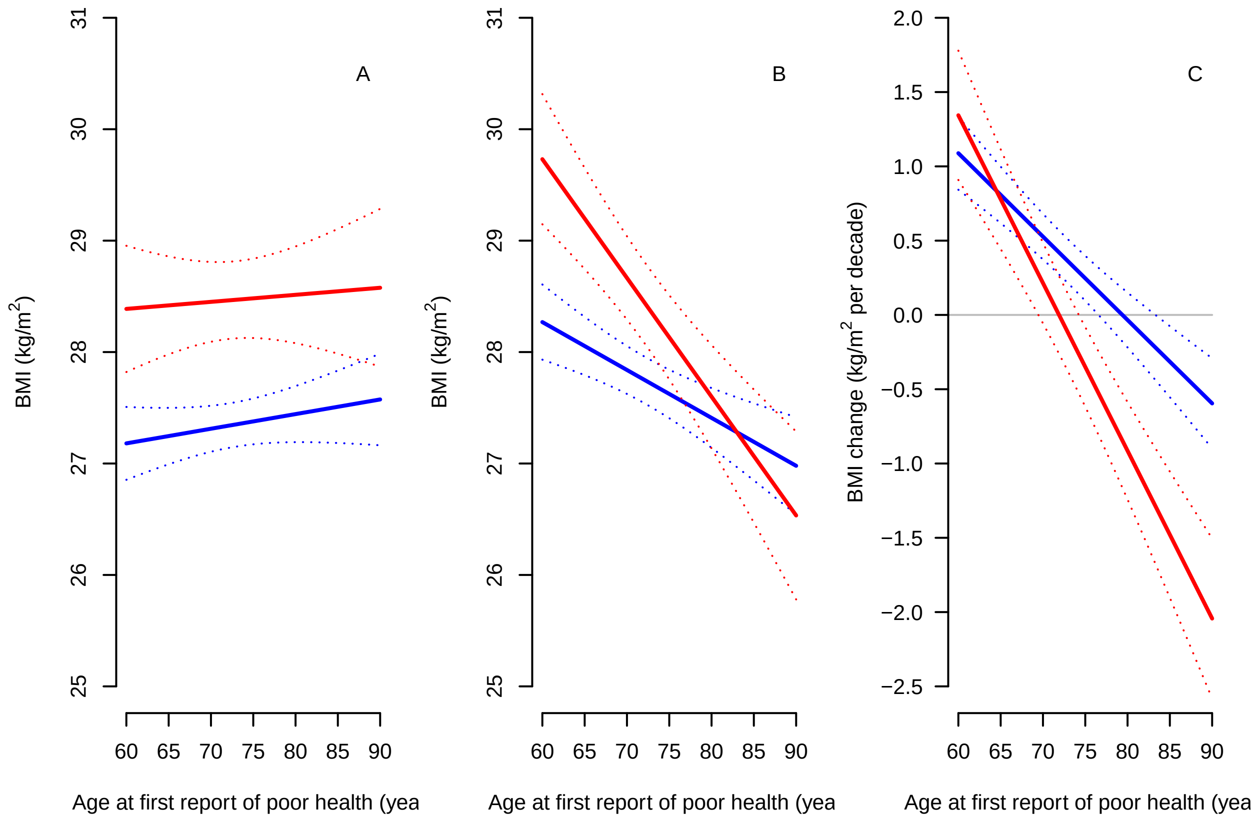


```
par(mfrow=c(3,2))
mainPlot()
smokePlot()
dmPlot()
cvdPlot()
cancerPlot()
lungPlot()
```

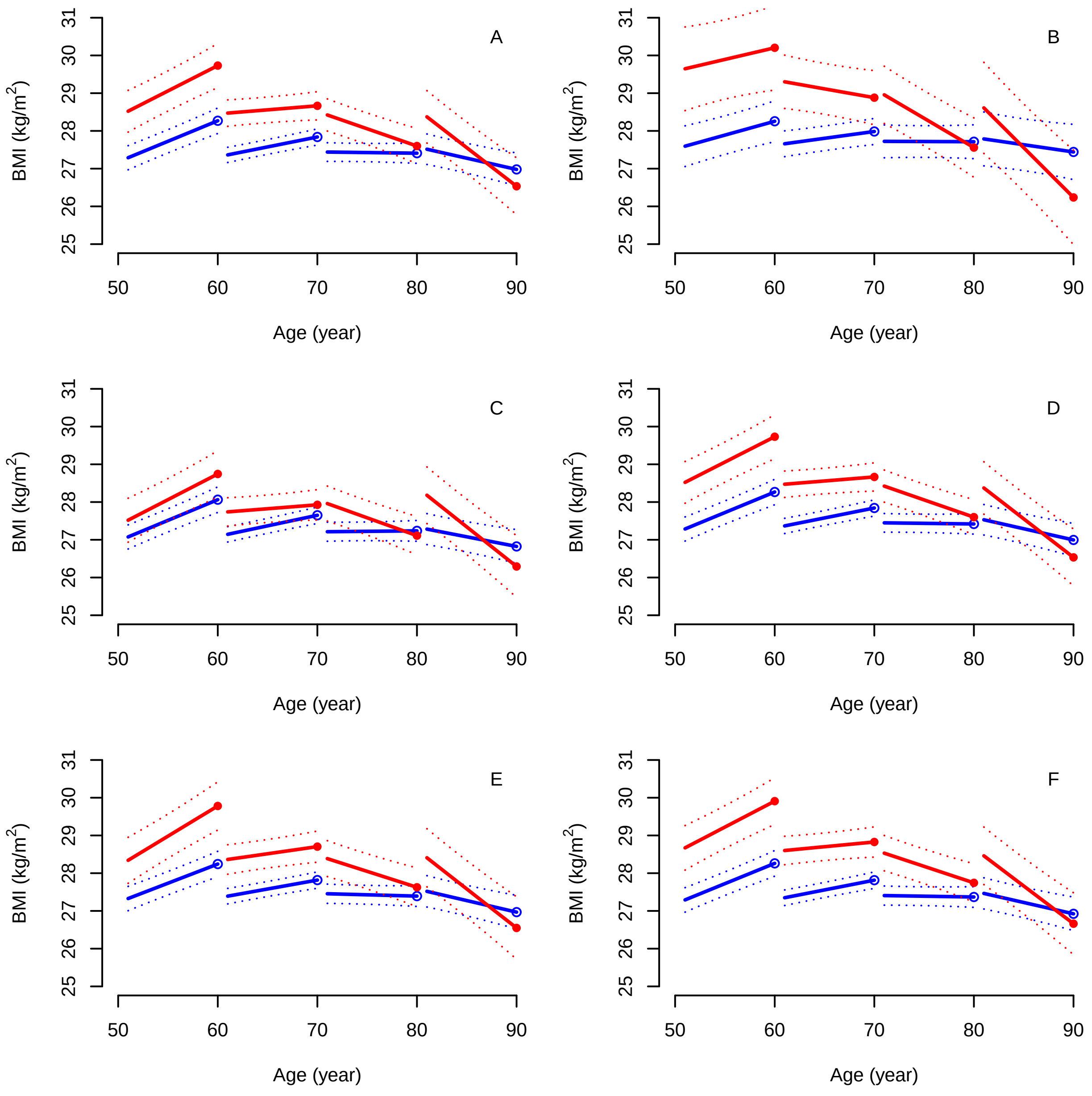


### Export figures in eps format

```
setEPS()
postscript("~/Google Drive/Research_AU/ELSA_section/fig2a.eps", height = 4, width = 4)
levelPlot()

postscript("~/Google Drive/Research_AU/ELSA_section/fig2b.eps", height = 4, width = 4)
levelPlot10y()

postscript("~/Google Drive/Research_AU/ELSA_section/fig2c.eps", height = 4, width = 4)
slopePlot()

postscript("~/Google Drive/Research_AU/ELSA_section/fig3a.eps", height = 4, width = 4)
```

```r
mainPlot()

postscript("~/Google Drive/Research_AU/ELSA_section/fig3c.eps", height = 4, width = 4)
dmPlot()

postscript("~/Google Drive/Research_AU/ELSA_section/fig3f.eps", height = 4, width = 4)
lungPlot()

postscript("~/Google Drive/Research_AU/ELSA_section/fig3b.eps", height = 4, width = 4)
smokePlot()

postscript("~/Google Drive/Research_AU/ELSA_section/fig3e.eps", height = 4, width = 4)
cancerPlot()

postscript("~/Google Drive/Research_AU/ELSA_section/fig3d.eps", height = 4, width = 4)
cvdPlot()

dev.off()
```

```
## pdf
##   2
```

### Variance components

In this section, we calculate the proportion of between-person variation explained by age at diagnosis. As the focus is on the variance components, we fit the models using REML instead of ML.

```r
modelA <- lme(bmi ~ t*case,
              data = longFinal,
              random=~t|id,
              na.action = na.omit,
              method="REML",
              control = lmeControl(msMaxIter = 250, opt="optim"))

VarCorr(modelA)
```

```
## id = pdLogChol(t)
##             Variance    StdDev    Corr
## (Intercept) 23.30760410 4.8277950 (Intr)
## t            0.04148874 0.2036878 0.443
## Residual     1.63412967 1.2783308
```

```r
modelB <- lme(bmi ~ t*diagAge*case,
              data = longFinal,
              random=~t|id,
              na.action = na.omit,
              method="REML",
              control = lmeControl(msMaxIter = 250, opt="optim"))

VarCorr(modelB)
```

```
## id = pdLogChol(t)
##             Variance    StdDev   Corr
## (Intercept) 22.95634567 4.791278 (Intr)
## t            0.03609695 0.189992 0.432
```

```
## Residual      1.63993900 1.280601
```

```r
# variance components (how age changes everything...)
rint <- as.numeric(c(VarCorr(modelA)[1], VarCorr(modelB)[1]))
rslope <- as.numeric(c(VarCorr(modelA)[2], VarCorr(modelB)[2]))

100*(rint[1]-rint[2])/rint[1]
```

```
## [1] 1.507055
```

```r
100*(rslope[1]-rslope[2])/rslope[1]
```

```
## [1] 12.99579
```

### Three-way interactions

Three-way interactions from the main (unadjusted) and the diabetes model.

```r
colIndex2 <- c(1,5,6,4)

# all
kable(ci.lin(modelFinal)[-1, colIndex2], "latex", booktabs = T) %>%
    kable_styling(latex_options = c("striped", "hold_position", "repeat_header"))
```

| | Estimate | 2.5% | 97.5% | P |
|---|---|---|---|---|
| t | 0.4456355 | 0.3325526 | 0.5587185 | 0.0000000 |
| diagAge | -0.0429938 | -0.0645807 | -0.0214069 | 0.0000948 |
| casecase | 5.2771892 | 2.1399029 | 8.4144755 | 0.0009778 |
| t:diagAge | -0.0056130 | -0.0071536 | -0.0040725 | 0.0000000 |
| t:casecase | 0.3661179 | 0.1358646 | 0.5963711 | 0.0018303 |
| diagAge:casecase | -0.0635870 | -0.1066076 | -0.0205665 | 0.0037681 |
| t:diagAge:casecase | -0.0056768 | -0.0088147 | -0.0025388 | 0.0003916 |

```r
# diabetes
kable(ci.lin(modelDM)[-1, colIndex2], "latex", booktabs = T) %>%
    kable_styling(latex_options = c("striped", "hold_position", "repeat_header"))
```

| | Estimate | 2.5% | 97.5% | P |
|---|---|---|---|---|
| t | 0.4310669 | 0.3148253 | 0.5473085 | 0.0000000 |
| diagAge | -0.0413110 | -0.0631244 | -0.0194975 | 0.0002058 |
| casecase | 3.1050210 | -0.1854373 | 6.3954792 | 0.0643845 |
| t:diagAge | -0.0053567 | -0.0069442 | -0.0037693 | 0.0000000 |
| t:casecase | 0.3967708 | 0.1506762 | 0.6428653 | 0.0015777 |
| diagAge:casecase | -0.0404004 | -0.0857083 | 0.0049076 | 0.0805213 |
| t:diagAge:casecase | -0.0061722 | -0.0095351 | -0.0028094 | 0.0003215 |

### Age when trajectories start decline

```r
# reminder of an earlier defined variable: ageAtDiag <- seq(60, 90, 1)
```

```
# by age at diagnosis
caseLevelMatrix <- cbind(1,0,ageAtDiag,1,0,0,ageAtDiag,0)
controlLevelMatrix <- cbind(1,0,ageAtDiag,0,0,0,0,0)
caseSlopeMatrix <- cbind(0,1,0,0,ageAtDiag,1,0,ageAtDiag)
controlSlopeMatrix <- cbind(0,1,0,0,ageAtDiag,0,0,0)

ctrLevelMatrix <- caseLevelMatrix - controlLevelMatrix
ctrSlopeMatrix <- caseSlopeMatrix - controlSlopeMatrix

# by age at diagnosis adjusted for SES
caseLevelSESMatrix <- cbind(1,0,ageAtDiag,1,0,0,0,0,0,ageAtDiag,0)
controlLevelSESMatrix <- cbind(1,0,ageAtDiag,0,0,0,0,0,0,0,0)
caseSlopeSESMatrix <- cbind(0,1,0,0,0,0,0,ageAtDiag,1,0,ageAtDiag)
controlSlopeSESMatrix <- cbind(0,1,0,0,0,0,0,ageAtDiag,0,0,0)

ctrLevelSESMatrix <- caseLevelSESMatrix - controlLevelSESMatrix
ctrSlopeSESMatrix <- caseSlopeSESMatrix - controlSlopeSESMatrix


# level
caseLevel <- round(cbind(ageAtDiag,
                        ci.lin(modelFinal, ctr.mat = caseLevelMatrix)[,colIndex],
                        ci.lin(modelSES, ctr.mat = caseLevelSESMatrix)[,colIndex]), 2)

controlLevel <- round(cbind(ageAtDiag,
                           ci.lin(modelFinal, ctr.mat = controlLevelMatrix)[,colIndex],
                           ci.lin(modelSES, ctr.mat=controlLevelSESMatrix)[,colIndex]),2)

levelDiff <- round(cbind(ageAtDiag,
                        ci.lin(modelFinal, ctr.mat = ctrLevelMatrix)[,colIndex],
                        ci.lin(modelSES, ctr.mat = ctrLevelSESMatrix)[,colIndex]), 2)

# slope
caseSlope <- round(cbind(ageAtDiag,
                        10*ci.lin(modelFinal, ctr.mat = caseSlopeMatrix)[,colIndex],
                        10*ci.lin(modelSES, ctr.mat = caseSlopeSESMatrix)[,colIndex]), 2)

controlSlope <- round(cbind(ageAtDiag,
                           10*ci.lin(modelFinal, ctr.mat=controlSlopeMatrix)[,colIndex],
                           10*ci.lin(modelSES, ctr.mat=controlSlopeSESMatrix)[,colIndex]), 2)

slopeDiff <- round(cbind(ageAtDiag,
                        10*ci.lin(modelFinal, ctr.mat = ctrSlopeMatrix)[,colIndex],
                        10*ci.lin(modelSES, ctr.mat = ctrSlopeSESMatrix)[,colIndex]), 2)

# level 10 years before
caseLevelMatrix10 <- cbind(1,-10,ageAtDiag,1,-10*ageAtDiag,-10,ageAtDiag,-10*ageAtDiag) # cases
controlLevelMatrix10 <- cbind(1,-10,ageAtDiag,0,-10*ageAtDiag,0,0,0) # controls
ctrLevelMatrix10 <- caseLevelMatrix10 - controlLevelMatrix10 # difference

level10ybefore <- round(cbind(ageAtDiag,
                        ci.lin(modelFinal, ctr.mat = caseLevelMatrix10)[,colIndex],
                        ci.lin(modelFinal, ctr.mat = controlLevelMatrix10)[,colIndex],
```

```r
                         ci.lin(modelFinal, ctr.mat = ctrLevelMatrix10)[,colIndex]), 2)

#summary(modelFinal)
```

```r
latopt <- c("striped", "hold_position", "scale_down", "repeat_header")
# LEVEL
# unadjusted
levelTable <- cbind(caseLevel[,1:4], controlLevel[,2:4], levelDiff[,2:4])

kable(levelTable, "latex", booktabs = T, caption = "BMI levels by age at first report of
      poor health") %>%
    kable_styling(latex_options = latopt) %>%
    add_header_above(c(" " = 1, "Case" = 3, "Control" = 3, "Diff." = 3))
```

```r
levelTableSES <- cbind(caseLevel[,c(1, 5:7)], controlLevel[,5:7], levelDiff[,5:7])

kable(levelTableSES, "latex", booktabs = T, caption = "SES-adjusted BMI levels by age at
      first report of poor health") %>%
    kable_styling(latex_options = latopt) %>%
    add_header_above(c(" " = 1, "Case" = 3, "Control" = 3, "Diff." = 3))
```

```r
# SLOPE (per decade)
slopeTable <- cbind(caseSlope[,1:4], controlSlope[,2:4], slopeDiff[,2:4])

kable(slopeTable, "latex", booktabs = T, caption = "BMI slopes (per decade) by age at
      first report of poor health") %>%
    kable_styling(latex_options = latopt) %>%
    add_header_above(c(" " = 1, "Case" = 3, "Control" = 3, "Diff." = 3))
```

```r
slopeTableSES <- cbind(caseSlope[,c(1, 5:7)], controlSlope[,5:7], slopeDiff[,5:7])

kable(slopeTableSES, "latex", booktabs = T, caption = "SES-adjusted BMI slopes
      (per decade) by age at first report of poor health") %>%
    kable_styling(latex_options = latopt) %>%
    add_header_above(c(" " = 1, "Case" = 3, "Control" = 3, "Diff." = 3))
```

```r
# 10-y before, unadjusted
kable(level10ybefore, "latex", booktabs = T, caption = "BMI levels 10 yrs before first report of
      poor health by age at first report") %>%
    kable_styling(latex_options = latopt) %>%
    add_header_above(c(" " = 1, "Case" = 3, "Control" = 3, "Diff." = 3))
```

BMI levels by age at first report of poor health

| | Case | | | Control | | | Diff. | | |
|---|---|---|---|---|---|---|---|---|---|
| ageAtDiag | Estimate | 2.5% | 97.5% | Estimate | 2.5% | 97.5% | Estimate | 2.5% | 97.5% |
| 60 | 29.73 | 29.15 | 30.32 | 28.27 | 27.93 | 28.61 | 1.46 | 0.79 | 2.14 |
| 61 | 29.62 | 29.07 | 30.18 | 28.23 | 27.91 | 28.55 | 1.40 | 0.76 | 2.04 |
| 62 | 29.52 | 28.99 | 30.05 | 28.18 | 27.88 | 28.49 | 1.33 | 0.73 | 1.94 |
| 63 | 29.41 | 28.91 | 29.91 | 28.14 | 27.85 | 28.43 | 1.27 | 0.69 | 1.85 |
| 64 | 29.30 | 28.83 | 29.78 | 28.10 | 27.82 | 28.37 | 1.21 | 0.66 | 1.76 |
| 65 | 29.20 | 28.74 | 29.65 | 28.05 | 27.79 | 28.32 | 1.14 | 0.62 | 1.67 |
| 66 | 29.09 | 28.66 | 29.52 | 28.01 | 27.76 | 28.26 | 1.08 | 0.58 | 1.58 |
| 67 | 28.99 | 28.57 | 29.40 | 27.97 | 27.73 | 28.21 | 1.02 | 0.54 | 1.49 |
| 68 | 28.88 | 28.48 | 29.28 | 27.93 | 27.70 | 28.15 | 0.95 | 0.49 | 1.41 |
| 69 | 28.77 | 28.39 | 29.16 | 27.88 | 27.66 | 28.10 | 0.89 | 0.45 | 1.33 |
| 70 | 28.67 | 28.29 | 29.04 | 27.84 | 27.62 | 28.05 | 0.83 | 0.39 | 1.26 |
| 71 | 28.56 | 28.19 | 28.93 | 27.80 | 27.59 | 28.01 | 0.76 | 0.34 | 1.19 |
| 72 | 28.45 | 28.09 | 28.82 | 27.75 | 27.54 | 27.96 | 0.70 | 0.28 | 1.12 |
| 73 | 28.35 | 27.98 | 28.71 | 27.71 | 27.50 | 27.92 | 0.64 | 0.21 | 1.06 |
| 74 | 28.24 | 27.87 | 28.61 | 27.67 | 27.45 | 27.88 | 0.57 | 0.15 | 1.00 |
| 75 | 28.13 | 27.75 | 28.51 | 27.62 | 27.41 | 27.84 | 0.51 | 0.07 | 0.94 |
| 76 | 28.03 | 27.64 | 28.42 | 27.58 | 27.36 | 27.81 | 0.44 | 0.00 | 0.89 |
| 77 | 27.92 | 27.52 | 28.32 | 27.54 | 27.31 | 27.77 | 0.38 | -0.09 | 0.85 |
| 78 | 27.81 | 27.39 | 28.23 | 27.50 | 27.25 | 27.74 | 0.32 | -0.17 | 0.80 |
| 79 | 27.71 | 27.26 | 28.15 | 27.45 | 27.20 | 27.71 | 0.25 | -0.26 | 0.76 |
| 80 | 27.60 | 27.14 | 28.06 | 27.41 | 27.14 | 27.68 | 0.19 | -0.35 | 0.73 |
| 81 | 27.49 | 27.01 | 27.98 | 27.37 | 27.08 | 27.65 | 0.13 | -0.44 | 0.69 |
| 82 | 27.39 | 26.87 | 27.90 | 27.32 | 27.03 | 27.62 | 0.06 | -0.53 | 0.66 |
| 83 | 27.28 | 26.74 | 27.82 | 27.28 | 26.97 | 27.59 | 0.00 | -0.62 | 0.62 |
| 84 | 27.17 | 26.60 | 27.74 | 27.24 | 26.91 | 27.57 | -0.06 | -0.72 | 0.59 |
| 85 | 27.07 | 26.47 | 27.66 | 27.19 | 26.85 | 27.54 | -0.13 | -0.82 | 0.56 |
| 86 | 26.96 | 26.33 | 27.59 | 27.15 | 26.79 | 27.51 | -0.19 | -0.92 | 0.53 |
| 87 | 26.85 | 26.20 | 27.51 | 27.11 | 26.73 | 27.49 | -0.25 | -1.02 | 0.51 |
| 88 | 26.75 | 26.06 | 27.44 | 27.07 | 26.67 | 27.46 | -0.32 | -1.11 | 0.48 |
| 89 | 26.64 | 25.92 | 27.36 | 27.02 | 26.60 | 27.44 | -0.38 | -1.22 | 0.45 |
| 90 | 26.53 | 25.78 | 27.29 | 26.98 | 26.54 | 27.42 | -0.45 | -1.32 | 0.43 |

SES-adjusted BMI levels by age at first report of poor health

| | Case | | | Control | | | Diff. | | |
|---|---|---|---|---|---|---|---|---|---|
| ageAtDiag | Estimate | 2.5% | 97.5% | Estimate | 2.5% | 97.5% | Estimate | 2.5% | 97.5% |
| 60 | 29.29 | 28.65 | 29.93 | 28.00 | 27.61 | 28.39 | 1.29 | 0.61 | 1.97 |
| 61 | 29.20 | 28.58 | 29.81 | 27.97 | 27.59 | 28.34 | 1.23 | 0.58 | 1.88 |
| 62 | 29.10 | 28.51 | 29.69 | 27.93 | 27.56 | 28.30 | 1.17 | 0.55 | 1.78 |
| 63 | 29.00 | 28.44 | 29.56 | 27.89 | 27.54 | 28.25 | 1.11 | 0.52 | 1.69 |
| 64 | 28.90 | 28.36 | 29.44 | 27.86 | 27.51 | 28.20 | 1.04 | 0.49 | 1.60 |
| 65 | 28.80 | 28.28 | 29.32 | 27.82 | 27.49 | 28.16 | 0.98 | 0.45 | 1.51 |
| 66 | 28.71 | 28.21 | 29.21 | 27.79 | 27.46 | 28.11 | 0.92 | 0.42 | 1.42 |
| 67 | 28.61 | 28.12 | 29.09 | 27.75 | 27.43 | 28.07 | 0.86 | 0.38 | 1.34 |
| 68 | 28.51 | 28.04 | 28.98 | 27.72 | 27.40 | 28.03 | 0.80 | 0.33 | 1.26 |
| 69 | 28.41 | 27.96 | 28.87 | 27.68 | 27.37 | 27.98 | 0.73 | 0.29 | 1.18 |
| 70 | 28.32 | 27.87 | 28.76 | 27.64 | 27.34 | 27.95 | 0.67 | 0.24 | 1.11 |
| 71 | 28.22 | 27.78 | 28.66 | 27.61 | 27.31 | 27.91 | 0.61 | 0.18 | 1.04 |
| 72 | 28.12 | 27.68 | 28.56 | 27.57 | 27.27 | 27.87 | 0.55 | 0.12 | 0.97 |
| 73 | 28.02 | 27.58 | 28.46 | 27.54 | 27.24 | 27.84 | 0.49 | 0.06 | 0.91 |
| 74 | 27.92 | 27.48 | 28.37 | 27.50 | 27.20 | 27.81 | 0.42 | -0.01 | 0.85 |
| 75 | 27.83 | 27.38 | 28.28 | 27.47 | 27.16 | 27.77 | 0.36 | -0.08 | 0.80 |
| 76 | 27.73 | 27.27 | 28.19 | 27.43 | 27.11 | 27.74 | 0.30 | -0.16 | 0.75 |
| 77 | 27.63 | 27.16 | 28.10 | 27.39 | 27.07 | 27.72 | 0.24 | -0.24 | 0.71 |
| 78 | 27.53 | 27.05 | 28.02 | 27.36 | 27.03 | 27.69 | 0.17 | -0.32 | 0.67 |
| 79 | 27.43 | 26.93 | 27.94 | 27.32 | 26.98 | 27.66 | 0.11 | -0.40 | 0.63 |
| 80 | 27.34 | 26.81 | 27.86 | 27.29 | 26.94 | 27.64 | 0.05 | -0.49 | 0.59 |
| 81 | 27.24 | 26.70 | 27.78 | 27.25 | 26.89 | 27.61 | -0.01 | -0.58 | 0.56 |
| 82 | 27.14 | 26.57 | 27.71 | 27.22 | 26.84 | 27.59 | -0.07 | -0.68 | 0.53 |
| 83 | 27.04 | 26.45 | 27.64 | 27.18 | 26.79 | 27.57 | -0.14 | -0.77 | 0.50 |
| 84 | 26.95 | 26.33 | 27.56 | 27.14 | 26.74 | 27.55 | -0.20 | -0.86 | 0.47 |
| 85 | 26.85 | 26.20 | 27.49 | 27.11 | 26.69 | 27.53 | -0.26 | -0.96 | 0.44 |
| 86 | 26.75 | 26.08 | 27.42 | 27.07 | 26.64 | 27.51 | -0.32 | -1.06 | 0.41 |
| 87 | 26.65 | 25.95 | 27.35 | 27.04 | 26.59 | 27.49 | -0.38 | -1.15 | 0.39 |
| 88 | 26.55 | 25.82 | 27.29 | 27.00 | 26.53 | 27.47 | -0.45 | -1.25 | 0.36 |
| 89 | 26.46 | 25.70 | 27.22 | 26.97 | 26.48 | 27.45 | -0.51 | -1.35 | 0.34 |
| 90 | 26.36 | 25.57 | 27.15 | 26.93 | 26.43 | 27.43 | -0.57 | -1.45 | 0.31 |

BMI slopes (per decade) by age at first report of poor health

| | Case | | | Control | | | Diff. | | |
|---|---|---|---|---|---|---|---|---|---|
| ageAtDiag | Estimate | 2.5% | 97.5% | Estimate | 2.5% | 97.5% | Estimate | 2.5% | 97.5% |
| 60 | 1.34 | 0.91 | 1.78 | 1.09 | 0.84 | 1.33 | 0.26 | -0.24 | 0.75 |
| 61 | 1.23 | 0.82 | 1.64 | 1.03 | 0.80 | 1.27 | 0.20 | -0.28 | 0.67 |
| 62 | 1.12 | 0.73 | 1.51 | 0.98 | 0.75 | 1.20 | 0.14 | -0.31 | 0.59 |
| 63 | 1.00 | 0.63 | 1.38 | 0.92 | 0.71 | 1.13 | 0.08 | -0.34 | 0.51 |
| 64 | 0.89 | 0.54 | 1.25 | 0.86 | 0.66 | 1.06 | 0.03 | -0.38 | 0.43 |
| 65 | 0.78 | 0.44 | 1.11 | 0.81 | 0.62 | 1.00 | -0.03 | -0.41 | 0.36 |
| 66 | 0.67 | 0.35 | 0.99 | 0.75 | 0.57 | 0.93 | -0.09 | -0.45 | 0.28 |
| 67 | 0.55 | 0.25 | 0.86 | 0.70 | 0.52 | 0.87 | -0.14 | -0.49 | 0.21 |
| 68 | 0.44 | 0.15 | 0.73 | 0.64 | 0.47 | 0.80 | -0.20 | -0.53 | 0.14 |
| 69 | 0.33 | 0.05 | 0.61 | 0.58 | 0.42 | 0.74 | -0.26 | -0.58 | 0.07 |
| 70 | 0.21 | -0.06 | 0.49 | 0.53 | 0.37 | 0.68 | -0.31 | -0.62 | 0.00 |
| 71 | 0.10 | -0.16 | 0.37 | 0.47 | 0.32 | 0.62 | -0.37 | -0.67 | -0.07 |
| 72 | -0.01 | -0.27 | 0.25 | 0.41 | 0.27 | 0.56 | -0.43 | -0.73 | -0.13 |
| 73 | -0.12 | -0.38 | 0.14 | 0.36 | 0.21 | 0.51 | -0.48 | -0.78 | -0.18 |
| 74 | -0.24 | -0.50 | 0.03 | 0.30 | 0.15 | 0.45 | -0.54 | -0.84 | -0.24 |
| 75 | -0.35 | -0.62 | -0.08 | 0.25 | 0.10 | 0.40 | -0.60 | -0.90 | -0.29 |
| 76 | -0.46 | -0.74 | -0.19 | 0.19 | 0.03 | 0.35 | -0.65 | -0.97 | -0.34 |
| 77 | -0.58 | -0.86 | -0.29 | 0.13 | -0.03 | 0.30 | -0.71 | -1.04 | -0.38 |
| 78 | -0.69 | -0.99 | -0.39 | 0.08 | -0.09 | 0.25 | -0.77 | -1.11 | -0.43 |
| 79 | -0.80 | -1.11 | -0.49 | 0.02 | -0.15 | 0.20 | -0.82 | -1.18 | -0.47 |
| 80 | -0.91 | -1.24 | -0.59 | -0.03 | -0.22 | 0.15 | -0.88 | -1.26 | -0.50 |
| 81 | -1.03 | -1.37 | -0.68 | -0.09 | -0.28 | 0.10 | -0.94 | -1.33 | -0.54 |
| 82 | -1.14 | -1.50 | -0.78 | -0.15 | -0.35 | 0.06 | -0.99 | -1.41 | -0.58 |
| 83 | -1.25 | -1.64 | -0.87 | -0.20 | -0.42 | 0.01 | -1.05 | -1.49 | -0.61 |
| 84 | -1.37 | -1.77 | -0.96 | -0.26 | -0.49 | -0.03 | -1.11 | -1.57 | -0.64 |
| 85 | -1.48 | -1.90 | -1.05 | -0.31 | -0.55 | -0.08 | -1.16 | -1.65 | -0.68 |
| 86 | -1.59 | -2.04 | -1.15 | -0.37 | -0.62 | -0.12 | -1.22 | -1.73 | -0.71 |
| 87 | -1.70 | -2.17 | -1.24 | -0.43 | -0.69 | -0.16 | -1.28 | -1.82 | -0.74 |
| 88 | -1.82 | -2.31 | -1.33 | -0.48 | -0.76 | -0.21 | -1.33 | -1.90 | -0.77 |
| 89 | -1.93 | -2.45 | -1.42 | -0.54 | -0.83 | -0.25 | -1.39 | -1.98 | -0.80 |
| 90 | -2.04 | -2.58 | -1.50 | -0.60 | -0.90 | -0.29 | -1.45 | -2.07 | -0.83 |

SES-adjusted BMI slopes (per decade) by age at first report of poor health

| | Case | | | Control | | | Diff. | | |
|---|---|---|---|---|---|---|---|---|---|
| ageAtDiag | Estimate | 2.5% | 97.5% | Estimate | 2.5% | 97.5% | Estimate | 2.5% | 97.5% |
| 60 | 1.27 | 0.83 | 1.71 | 1.09 | 0.84 | 1.34 | 0.18 | -0.33 | 0.69 |
| 61 | 1.17 | 0.75 | 1.59 | 1.03 | 0.80 | 1.27 | 0.13 | -0.35 | 0.61 |
| 62 | 1.06 | 0.66 | 1.46 | 0.98 | 0.76 | 1.20 | 0.08 | -0.37 | 0.54 |
| 63 | 0.96 | 0.58 | 1.33 | 0.92 | 0.71 | 1.14 | 0.03 | -0.40 | 0.47 |
| 64 | 0.85 | 0.49 | 1.21 | 0.87 | 0.67 | 1.07 | -0.02 | -0.43 | 0.40 |
| 65 | 0.75 | 0.41 | 1.09 | 0.81 | 0.62 | 1.00 | -0.07 | -0.46 | 0.33 |
| 66 | 0.64 | 0.32 | 0.96 | 0.76 | 0.57 | 0.94 | -0.11 | -0.49 | 0.26 |
| 67 | 0.54 | 0.23 | 0.84 | 0.70 | 0.53 | 0.87 | -0.16 | -0.52 | 0.19 |
| 68 | 0.43 | 0.13 | 0.73 | 0.64 | 0.48 | 0.81 | -0.21 | -0.55 | 0.13 |
| 69 | 0.32 | 0.04 | 0.61 | 0.59 | 0.43 | 0.75 | -0.26 | -0.59 | 0.06 |
| 70 | 0.22 | -0.06 | 0.49 | 0.53 | 0.38 | 0.69 | -0.31 | -0.63 | 0.00 |
| 71 | 0.11 | -0.16 | 0.38 | 0.48 | 0.32 | 0.63 | -0.36 | -0.67 | -0.05 |
| 72 | 0.01 | -0.26 | 0.27 | 0.42 | 0.27 | 0.57 | -0.41 | -0.72 | -0.11 |
| 73 | -0.10 | -0.36 | 0.17 | 0.36 | 0.21 | 0.51 | -0.46 | -0.76 | -0.16 |
| 74 | -0.20 | -0.47 | 0.07 | 0.31 | 0.16 | 0.46 | -0.51 | -0.82 | -0.20 |
| 75 | -0.31 | -0.58 | -0.03 | 0.25 | 0.10 | 0.41 | -0.56 | -0.87 | -0.25 |
| 76 | -0.41 | -0.69 | -0.13 | 0.20 | 0.04 | 0.35 | -0.61 | -0.93 | -0.29 |
| 77 | -0.52 | -0.81 | -0.23 | 0.14 | -0.02 | 0.30 | -0.66 | -0.99 | -0.32 |
| 78 | -0.62 | -0.93 | -0.32 | 0.08 | -0.09 | 0.26 | -0.71 | -1.06 | -0.36 |
| 79 | -0.73 | -1.05 | -0.41 | 0.03 | -0.15 | 0.21 | -0.76 | -1.12 | -0.39 |
| 80 | -0.83 | -1.17 | -0.50 | -0.03 | -0.22 | 0.16 | -0.81 | -1.19 | -0.42 |
| 81 | -0.94 | -1.29 | -0.59 | -0.08 | -0.28 | 0.12 | -0.85 | -1.26 | -0.45 |
| 82 | -1.04 | -1.41 | -0.67 | -0.14 | -0.35 | 0.07 | -0.90 | -1.33 | -0.48 |
| 83 | -1.15 | -1.54 | -0.76 | -0.20 | -0.42 | 0.03 | -0.95 | -1.40 | -0.50 |
| 84 | -1.25 | -1.67 | -0.84 | -0.25 | -0.49 | -0.02 | -1.00 | -1.48 | -0.53 |
| 85 | -1.36 | -1.79 | -0.93 | -0.31 | -0.55 | -0.06 | -1.05 | -1.55 | -0.55 |
| 86 | -1.46 | -1.92 | -1.01 | -0.36 | -0.62 | -0.11 | -1.10 | -1.62 | -0.58 |
| 87 | -1.57 | -2.05 | -1.09 | -0.42 | -0.69 | -0.15 | -1.15 | -1.70 | -0.60 |
| 88 | -1.68 | -2.18 | -1.17 | -0.48 | -0.76 | -0.19 | -1.20 | -1.78 | -0.62 |
| 89 | -1.78 | -2.31 | -1.26 | -0.53 | -0.83 | -0.23 | -1.25 | -1.85 | -0.64 |
| 90 | -1.89 | -2.44 | -1.34 | -0.59 | -0.90 | -0.28 | -1.30 | -1.93 | -0.67 |

BMI levels 10 yrs before first report of poor health by age at first report

| | Case | | | Control | | | Diff. | | |
|---|---|---|---|---|---|---|---|---|---|
| ageAtDiag | Estimate | 2.5% | 97.5% | Estimate | 2.5% | 97.5% | Estimate | 2.5% | 97.5% |
| 60 | 28.39 | 27.82 | 28.95 | 27.18 | 26.85 | 27.51 | 1.21 | 0.55 | 1.86 |
| 61 | 28.39 | 27.86 | 28.93 | 27.19 | 26.88 | 27.50 | 1.20 | 0.58 | 1.82 |
| 62 | 28.40 | 27.89 | 28.91 | 27.21 | 26.91 | 27.50 | 1.19 | 0.60 | 1.78 |
| 63 | 28.41 | 27.92 | 28.89 | 27.22 | 26.94 | 27.50 | 1.19 | 0.63 | 1.75 |
| 64 | 28.41 | 27.95 | 28.87 | 27.23 | 26.97 | 27.50 | 1.18 | 0.65 | 1.71 |
| 65 | 28.42 | 27.98 | 28.86 | 27.25 | 26.99 | 27.50 | 1.17 | 0.67 | 1.68 |
| 66 | 28.43 | 28.01 | 28.84 | 27.26 | 27.02 | 27.50 | 1.17 | 0.68 | 1.65 |
| 67 | 28.43 | 28.03 | 28.83 | 27.27 | 27.04 | 27.50 | 1.16 | 0.70 | 1.62 |
| 68 | 28.44 | 28.06 | 28.82 | 27.29 | 27.07 | 27.51 | 1.15 | 0.71 | 1.59 |
| 69 | 28.44 | 28.08 | 28.81 | 27.30 | 27.09 | 27.51 | 1.15 | 0.72 | 1.57 |
| 70 | 28.45 | 28.09 | 28.81 | 27.31 | 27.11 | 27.52 | 1.14 | 0.73 | 1.55 |
| 71 | 28.46 | 28.11 | 28.81 | 27.33 | 27.12 | 27.53 | 1.13 | 0.73 | 1.54 |
| 72 | 28.46 | 28.12 | 28.81 | 27.34 | 27.14 | 27.54 | 1.13 | 0.73 | 1.52 |
| 73 | 28.47 | 28.12 | 28.82 | 27.35 | 27.15 | 27.55 | 1.12 | 0.72 | 1.52 |
| 74 | 28.48 | 28.13 | 28.83 | 27.36 | 27.16 | 27.57 | 1.11 | 0.71 | 1.51 |
| 75 | 28.48 | 28.13 | 28.84 | 27.38 | 27.17 | 27.58 | 1.10 | 0.69 | 1.52 |
| 76 | 28.49 | 28.12 | 28.85 | 27.39 | 27.18 | 27.60 | 1.10 | 0.67 | 1.52 |
| 77 | 28.49 | 28.12 | 28.87 | 27.40 | 27.18 | 27.62 | 1.09 | 0.65 | 1.53 |
| 78 | 28.50 | 28.11 | 28.90 | 27.42 | 27.19 | 27.65 | 1.08 | 0.63 | 1.54 |
| 79 | 28.51 | 28.09 | 28.92 | 27.43 | 27.19 | 27.67 | 1.08 | 0.60 | 1.56 |
| 80 | 28.51 | 28.08 | 28.95 | 27.44 | 27.19 | 27.69 | 1.07 | 0.57 | 1.57 |
| 81 | 28.52 | 28.06 | 28.98 | 27.46 | 27.19 | 27.72 | 1.06 | 0.54 | 1.59 |
| 82 | 28.53 | 28.05 | 29.01 | 27.47 | 27.19 | 27.75 | 1.06 | 0.50 | 1.61 |
| 83 | 28.53 | 28.03 | 29.04 | 27.48 | 27.19 | 27.78 | 1.05 | 0.47 | 1.63 |
| 84 | 28.54 | 28.01 | 29.07 | 27.50 | 27.19 | 27.80 | 1.04 | 0.43 | 1.66 |
| 85 | 28.55 | 27.99 | 29.11 | 27.51 | 27.18 | 27.83 | 1.04 | 0.39 | 1.68 |
| 86 | 28.55 | 27.96 | 29.14 | 27.52 | 27.18 | 27.86 | 1.03 | 0.35 | 1.71 |
| 87 | 28.56 | 27.94 | 29.18 | 27.54 | 27.18 | 27.89 | 1.02 | 0.31 | 1.74 |
| 88 | 28.56 | 27.92 | 29.21 | 27.55 | 27.17 | 27.92 | 1.02 | 0.27 | 1.76 |
| 89 | 28.57 | 27.89 | 29.25 | 27.56 | 27.17 | 27.95 | 1.01 | 0.23 | 1.79 |
| 90 | 28.58 | 27.87 | 29.29 | 27.57 | 27.16 | 27.99 | 1.00 | 0.18 | 1.82 |
